# A novel isoform of Tensin1 promotes actin filament assembly for efficient erythroblast enucleation

**DOI:** 10.1101/2024.12.13.628322

**Authors:** Arit Ghosh, Megan Coffin, Dimitri M. Diaz, Sarah Barndt, Vincent Schulz, Patrick Gallagher, Su Hao Lo, Velia M. Fowler

**Affiliations:** Department of Biological Sciences, University of Delaware, Newark, DE; Delaware Biotechnology Institute, UD Flow Cytometry Core, Newark, DE; Department of Medical and Molecular Sciences, University of Delaware, Newark, DE; Department of Pediatrics, Yale University, New Haven, CT; Department of Pediatrics, Ohio State University, Columbus, OH; Department of Biochemistry and Molecular Medicine, University of California-Davis, Sacramento, CA 95817, USA; Cancer and Immunology Research Center, National Yang Ming Chiao Tung University, Taipei 112304, Taiwan; Institute of Molecular and Genomic Medicine, National Health of Research Institutes, Miaoli 35053, Taiwan

**Keywords:** Actin Polymerization, Enucleation, Enucleosome, Erythroid Differentiation, eTNS1

## Abstract

Mammalian red blood cells are generated via a terminal erythroid differentiation pathway culminating in cell polarization and enucleation. Actin filament polymerization is critical for enucleation, but the molecular regulatory mechanisms remain poorly understood. We utilized publicly available RNA-seq and proteomics datasets to mine for actin-binding proteins and actin-nucleation factors differentially expressed during human erythroid differentiation and discovered that a focal adhesion protein—Tensin-1—dramatically increases in expression late in differentiation. Remarkably, we found that differentiating human CD34+ cells express a novel truncated form of Tensin-1 (eTNS1; Mr ∼125 kDa) missing the N-terminal half of the protein, due to an internal mRNA translation start site resulting in a unique exon 1. eTNS1 localized to the cytoplasm during terminal erythroid differentiation, with no apparent membrane association or focal adhesion formation. Knocking out eTNS1 had no effect on assembly of the spectrin membrane skeleton but led to impaired enucleation and absent or mis-localized actin filament foci in enucleating erythroblasts. We conclude that eTNS1 is a novel regulator of actin filament assembly during human erythroid terminal differentiation required for efficient enucleation.

## Introduction

Erythropoiesis in mammals is a complex process that begins with hematopoietic stem cell (HSC) commitment to the erythroid lineage, proliferation of erythroid progenitor cells, followed by terminal differentiation, with gradually decreasing cell and nuclear size, culminating in cell cycle exit, followed by enucleation to yield a reticulocyte and a pyrenocyte (expelled nucleus) (Chen et al., 2009; Ji et al., 2011; Keerthivasan et al., 2011; Newton et al., 2024; Simpson & Kling, 1967). During terminal differentiation, erythroblasts also undergo membrane remodeling, mitochondria and organelle loss, as well as cytoskeletal reorganization to form a two-dimensional spectrin-based membrane skeleton comprised of α1,β1-spectrin and short actin filaments (F-actin) along with other components (Fowler, 2013; Lux, 2016; Risinger & Kalfa, 2020). The unique qualities of the F-actin-spectrin membrane skeleton and absence of a nucleus confer mechanical stability and remarkable deformability properties on the mature biconcave erythrocytes, enabling them to efficiently navigate narrow microcapillaries and carry out oxygen and C02 transfer (Diez-Silva et al., 2010; Mohandas & Gallagher, 2008; Risinger & Kalfa, 2020). Several blood disorders, including hemolytic anemias, dyserythropoietic anemias, and myelodysplasias, have been linked to altered membrane deformability and stability that may result from impaired enucleation mechanisms (An et al., 2023; Buks et al., 2021; Iolascon et al., 2020; King et al., 2022; Mohandas & Gallagher, 2008; Ramos et al., 2013; Risinger et al., 2019; Risinger & Kalfa, 2020).

Enucleation comprises a series of dramatic morphological transformations that involve nuclear polarization to one side of the cell, membrane and organelle remodeling to remove unwanted components from the incipient reticulocyte and retain others, followed by nuclear translocation and extrusion, and finally, separation of the pyrenocyte from the reticulocyte (Ji et al., 2011; Moras et al., 2017; Newton et al., 2024; Palis, 2014). Both the actin cytoskeleton and microtubules are required for efficient enucleation, with microtubule assembly promoting nuclear polarization, while F-actin assembly and non-muscle myosin IIB (NMIIB) contractility are essential for nuclear extrusion (Kobayashi et al., 2016; Koury et al., 1989; Ubukawa et al., 2020; Ubukawa et al., 2012; Wang et al., 2012). In both mouse and human erythroblasts, immediately prior to nuclear expulsion, F-actin and NMIIB assemble into a prominent cytoplasmic structure at the rear of the nucleus, termed the enucleosome, which is proposed to provide forces to drive nuclear extrusion (Nowak et al., 2017). In mouse, but not human erythroblasts, F-actin and NMIIB also assemble into a ring-like structure surrounding the cellular constriction site, termed the contractile F-actin ring, which may also provide forces for extrusion (Konstantinidis et al., 2012; Koury et al., 1989; Wang et al., 2012). Although several actin-regulatory pathways, including signaling via Ca^++^-Calmodulin, Phosphoinositide-3 kinase (PI3K), Rac1/2, CDC42 and ROCK, are involved in enucleation, (Ji et al., 2008; Konstantinidis et al., 2012; Koury et al., 1989; Ubukawa et al., 2020; Ubukawa et al., 2012; Wang et al., 2012; Wölwer et al., 2016), the downstream effectors and actin-binding proteins regulating F-actin polymerization into these distinct specialized cytoskeletal structures during enucleation are not understood.

To investigate a role for actin-binding proteins (ABPs) during terminal erythroid differentiation and enucleation, we adopted a data-mining approach to identify well-known ABPs and actin-nucleation factors (NFs) (Appendix Table S1) within comprehensive transcriptomics (RNA-seq) and proteomics (LC-MS) datasets for stages of erythroid terminal differentiation (Gautier et al., 2016; Yan et al., 2018). We discovered that a single actin-binding protein, Tensin1 (TNS1), was highly upregulated transcriptionally and translationally late in human erythroid differentiation from CD34+ cells. TNS1 is a ∼220 kDa focal adhesion (FA) molecule that belongs to the tensin family of proteins, which are involved in cellular adhesion, polarization, migration, proliferation, and invasion (Blangy, 2017; Liao & Lo, 2021; Wang et al., 2022). TNS1 has a focal adhesion binding site (FAB-N) in the N-terminal region that overlaps with an actin-binding domain (ABD-I), a second actin-binding domain that is centrally located (ABD-II); as well as another focal adhesion binding site (FAB-C) in the C-terminal end that overlaps with Src homology 2 (SH2) and phosphotyrosine-binding (PTB) domains (Liao & Lo, 2021). The multi-domain structure allows TNS1 to localize to integrin-mediated focal adhesions (Chen & Lo, 2003; Lo, 2004) and act as an integrin adapter protein, linking the extracellular matrix to the actin cytoskeleton and signal transduction through numerous binding partners (Chen et al., 2002; Legate & Fässler, 2009; Liao & Lo, 2021).

In human erythroblasts, we discovered a novel short isoform of TNS1 (Mr ∼125kDa), termed erythroid tensin (eTNS1), that is expressed from an internal start site in a unique exon 1, which is present in humans and other primates, but not in rodents. The short eTNS1 protein is missing the N-terminal and internal actin-binding domains (ABD-I and II) but retains the FAB-C site. Surprisingly, eTNS1 localizes to the cytoplasm in polarized and enucleating erythroblasts, with no apparent membrane association or focal adhesion formation, and eTNS1 does not appear to colocalize with F-actin or be enriched in the enucleosome. Using confocal microscopy at a single cell resolution and image analyses we demonstrate that loss of eTNS1 nevertheless impairs F-actin assembly into the enucleosome in polarized and enucleating erythroblasts, and significantly reduces enucleation efficiency. Together, these observations identify eTNS1 as a novel regulator of F-actin assembly during human erythroid terminal differentiation and indicate that eTNS1 plays a critical role in promoting erythroblast enucleation.

## Results

### RNA-sequencing and proteomic database mining reveals increased expression of TNS1 during human terminal erythroid differentiation

Efficient enucleation depends on F-actin polymerization and reorganization into a prominent F-actin rich focus located at the rear of the translocating nucleus (enucleosome) (Figure 1A) (Nowak et al., 2017). To investigate the regulation of F-actin assembly into this structure during human erythroblast enucleation, we identified 135 known ABPs and NFs (Campellone & Welch, 2010; Chesarone & Goode, 2009; Fowler, 2013; Siton-Mendelson & Bernheim-Groswasser, 2017)(Appendix Table S1) and cross-referenced this list with publicly available RNA-sequencing and proteomic datasets to determine which of these ABPs/NFs increased during terminal erythroid differentiation (Figure 1B, Appendix Table S2) (Gautier et al., 2016; Yan et al., 2018). As a proof of concept, we conducted a similar analysis of 12 ABPs in the red cell membrane skeleton (Fowler, 2013; Lux, 2016) and 25 red cell transmembrane and membrane-associated proteins (Mohandas & Gallagher, 2008; Risinger & Kalfa, 2020) (Appendix Figure S1, Appendix Table S1). The fold change in expression was plotted on a linear scale (converted from log2 fold change) and normalized to the proerythroblast expression level for each hit (Figure 1C, D, Appendix Table S2). We found that TNS1 is highly upregulated at both the transcriptional (40-fold increase) and protein (30-fold increase) level in orthochromatic erythroblasts, compared to proerythroblasts (Figure 1C, D). TNS1 expression increased only in the late stages of erythroblast differentiation and was not enriched in proerythroblasts compared to BFU-E and CFU-E erythroid progenitor cells (Appendix Figure S2). This dramatic increase in TNS1 expression contrasts with other ABPs and NFs in our dataset that either decreased or showed a minimal increase in expression during terminal erythroid differentiation (Figure 1C, D). These results strongly suggest that TNS1 is involved in the late stages of human terminal erythroid differentiation.

**Figure 1:**
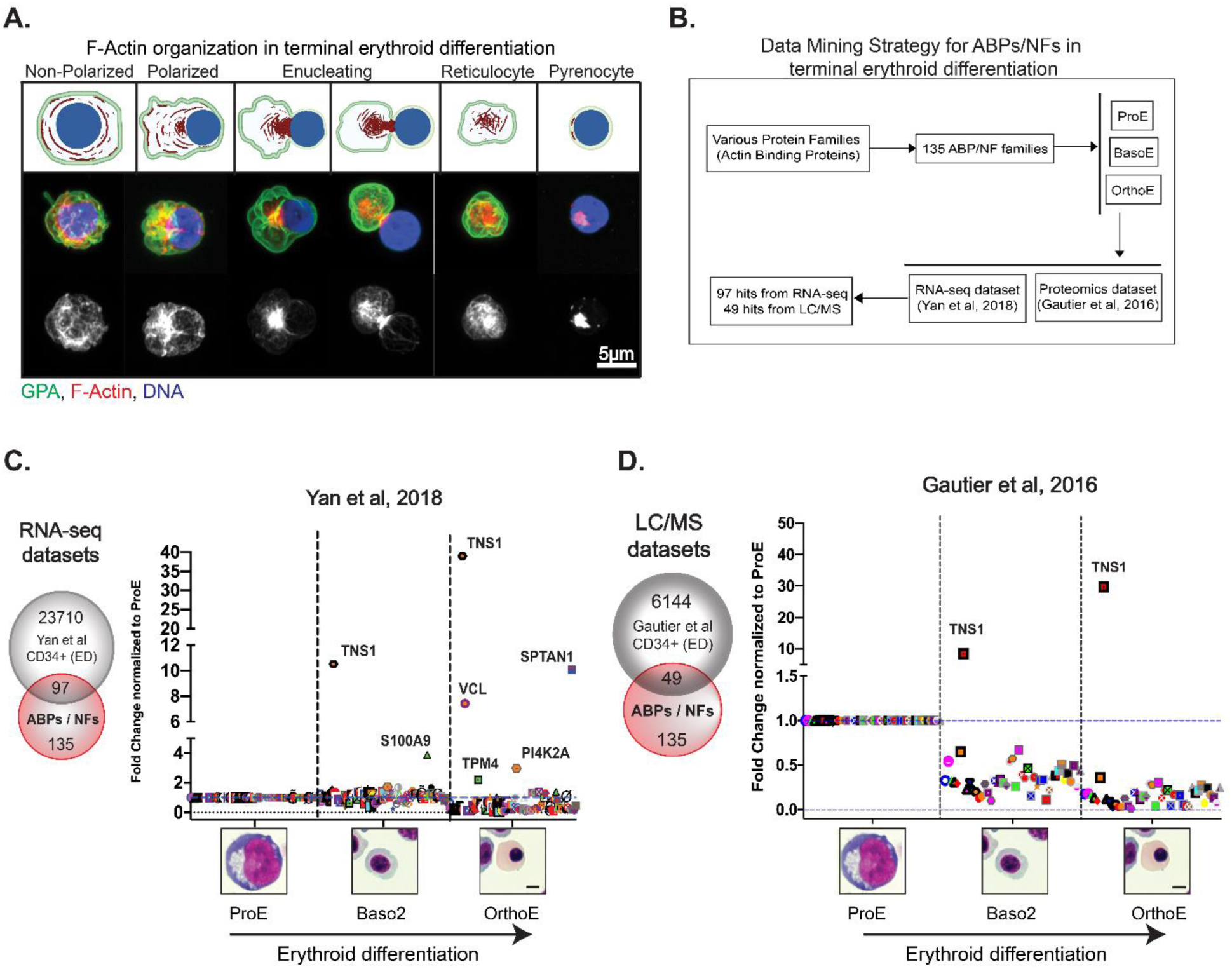
Database mining of actin-binding proteins and actin-nucleation factors reveals increased TNS1 expression during human terminal erythroid differentiation. (A-top) Schematic of F-actin reorganization (red) into the enucleosome, GPA sorting to the reticulocyte (green), and nuclear expulsion (blue) during human erythroblast enucleation, created with BioRender.com (A-middle) Maximum intensity projection of Airyscan Z-stacks of human CD34+ cells prior to and during enucleation immunostained for GPA (green), F-actin (phalloidin; red), and nuclei (Hoechst; blue). (A-bottom) F-actin staining in gray scale shows the formation of the enucleosome at the rear of the nucleus. Scale bar, 5µm. (B) Flowchart representing the data mining strategy for 135 actin-binding proteins (ABPs) and actin-nucleation factors (NFs) to identify mRNAs and proteins that are up-regulated during terminal erythroid differentiation of CD34+ cells. Graphs showing fold change in expression of (C) 97 RNA-sequencing and (D) 49 proteomics hits during terminal erythroid differentiation. Values were plotted on a linear scale (converted from log2 fold change) and normalized to their respective expression level at the proerythroblast stage. Representative Giemsa-stained cytospin images for the human erythroblast stages included in bioinformatics analysis (ProE = proerythroblast; Baso2 = basophilic erythroblast; OrthoE = orthochromatic erythroblast). Scale bar, 5µm.

### TNS1 expression increases during human terminal erythroid differentiation and enucleation in vitro

To investigate the role of TNS1 during erythropoiesis, we differentiated human cord blood-derived CD34+ cells towards the erythroid lineage using a three-phase culturing system (Dulmovits et al., 2016). Cells were collected from culture on days 7, 11, and 14 for molecular, biochemical, and morphological analysis. We confirmed CD34+ cells were differentiating along the erythroid pathway as expected by assessing changes in cell morphology using Giemsa staining (Appendix Figure S3) and by determining the expression of cell surface markers GPA, α4-integrin, and band 3 by flow cytometry (Appendix Figure S4A, C) (Hu et al., 2013). By day 14, Giemsa staining revealed a heterogeneous population of enucleating cells, reticulocytes, and pyrenocytes (Appendix Figure S3). DRAQ5 nuclear staining was used to differentiate between nucleated erythroblasts and enucleated reticulocytes to determine the extent of enucleation. On average, we found that 23% of cells had enucleated to become reticulocytes by day 14 of culture, and 38% had enucleated by day 17 of culture (Appendix Figure S4D).

To confirm the results from the bioinformatic analysis, we assessed *TNS1* mRNA expression at days 7, 11, and 14 in terminally differentiating erythroid cell cultures by TaqMan gene expression assay (Figure 2A). We used a TaqMan probe spanning exons 32-33 near the 3’ end of the human *TNS1* gene, which coincides with the PTB domain near the C-terminal end of the TNS1 protein (Figure 3B). qRT-PCR results confirmed a significant 12-fold increase in *TNS1* mRNA expression in day 14 cells, compared to day 7 cells (Figure 2A). Next, we examined TNS1 protein levels by Western blot using a polyclonal anti-peptide antibody to amino acids 1326-1339 of TNS1 and determined that TNS1 protein expression increased nearly 5-fold as CD34+ cells differentiated from day 7 to day 14 in culture (Figure 2B, C). qRT-PCR and Western blot analysis confirmed terminal erythroid differentiation based on an increase in erythroid membrane skeleton genes (SPTA1, EPB41) and proteins (α1β1-spectrin, protein 4.1R) that are upregulated during terminal erythroid differentiation (Figure 2A-C). Unlike U118 glioblastoma cells which express full-length TNS1 (Mr ∼220kDa), no full-length TNS1 is detected in erythroid cells, and instead, a prominent immunoreactive band at Mr ∼125kDa is present which increases from days 7 to 14 of culture (Figure 2B, C). This suggests that human erythroid cells may express a short isoform of TNS1 during late stages of terminal erythroid differentiation.

**Figure 2:**
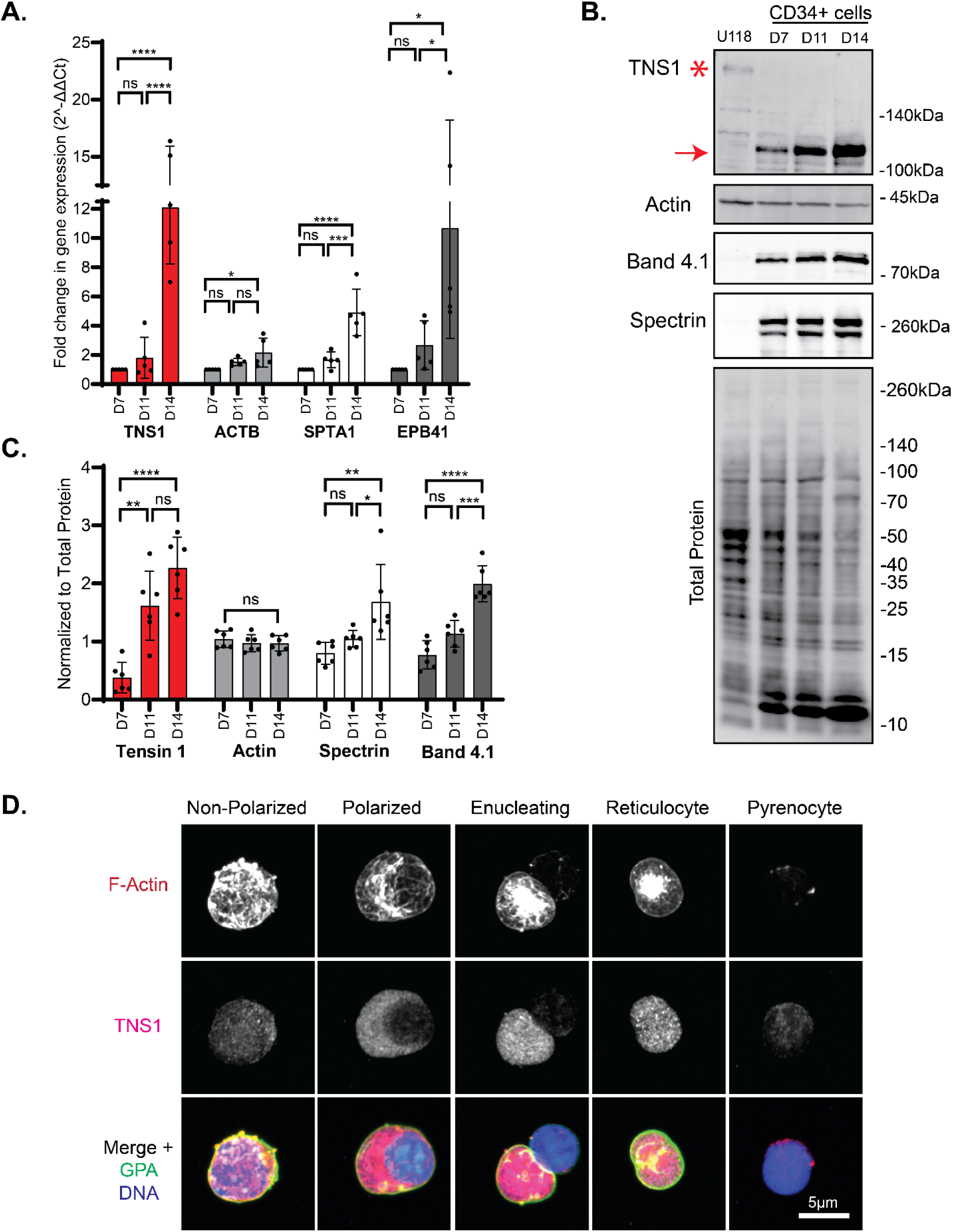
Expression of TNS1 is upregulated during in vitro human terminal erythroid differentiation. (A) mRNA levels of TNS1, β-actin (ACTB), α1-spectrin (SPTA1), and protein 4.1R (EPB41) in CD34+ cell cultures on days 7, 11, and 14 of differentiation, measured by TaqMan qRT-PCR. Values are mean ± SD from 5 individual CD34+ cultures. Fold-change in gene expression normalized to day 7 cells. *\*p<0.05; ***p<0.001; ****p<0.0001.* (B) Representative Western blots of TNS1, total actin, protein 4.1R, (α1β1)-spectrin, and total protein of differentiated CD34+ cells, and U118 glioblastoma cells; 15 µg protein loaded per lane. Asterisk, full-length TNS1 in U118 cells (Mr ∼220kDa); Red arrow, immunoreactive TNS1 polypeptide in erythroid cells (Mr ∼125kDa). (C) Quantification of immunoblots normalized to total protein. Band intensities analyzed by ImageJ. Values are mean ± SD from 6 individual CD34+ cultures. *\*p<0.05; **p<0.01; ***p<0.001; ****p<0.0001.* (D) Maximum intensity projections of Airyscan Z-stacks of cells from day 14 cultures, prior to and at various stages of enucleation, stained for F-actin (red), TNS1 (magenta), GPA (green), and nuclei (blue). Scale bar, 5µm.

**Figure 3.**
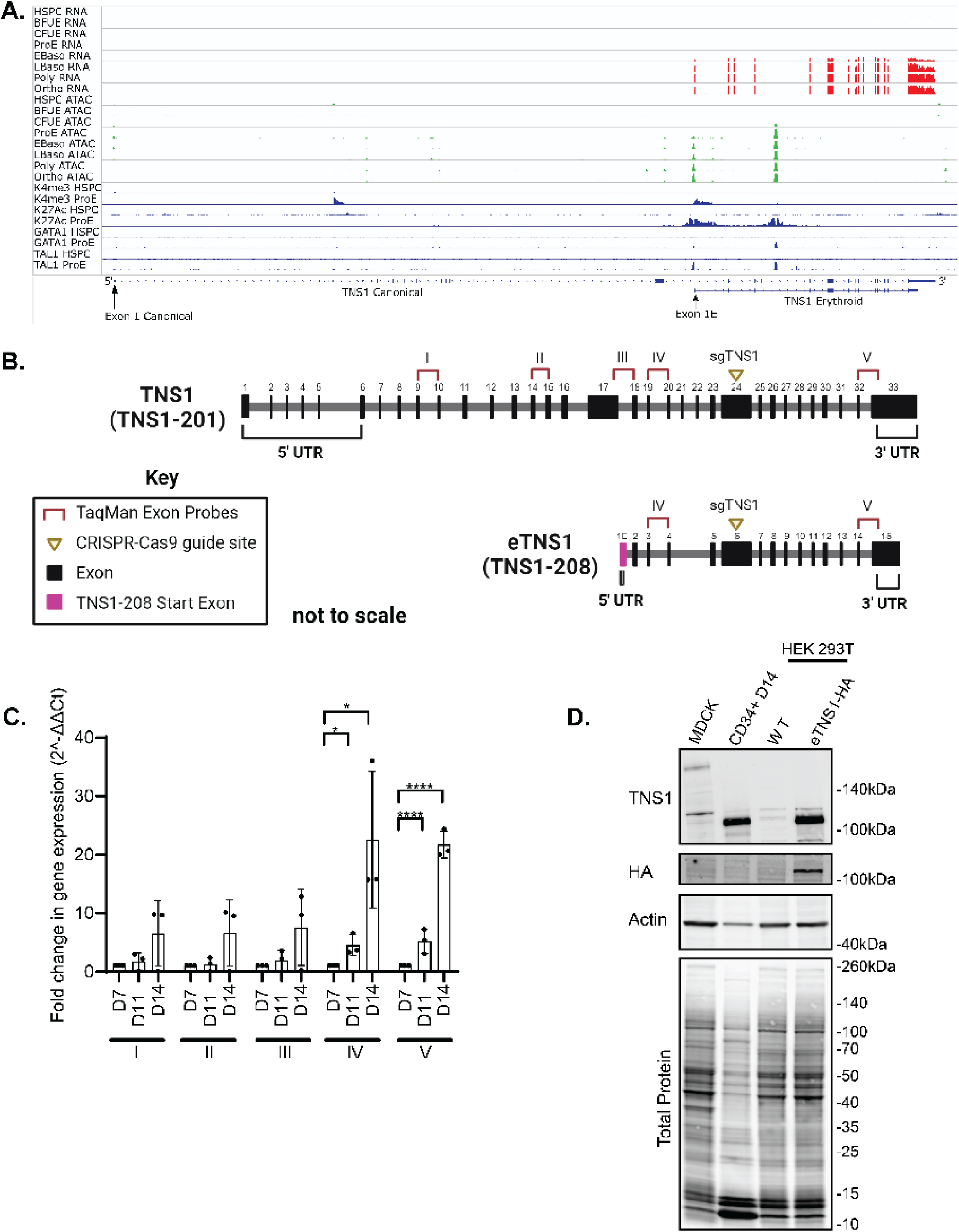
Human erythroblasts express a short isoform of TNS1 (eTNS1). (A) mRNA expression of TNS1 exons during human erythroid differentiation from human stem and progenitor cells (HSPC) through terminal differentiation to orthrochromatic erythroblasts (orthoE). Exons encoding the canonical *Ensembl TNS1* mRNA transcript and the erythroid *TNS1 (eTNS1)* mRNA transcript are shown at the bottom, with location of alternate exon 1s containing initiator methionines denoted by arrows. Top, red: mRNA expression, determined by RNA-seq, increases during terminal erythroid differentiation starting at the early basophilic erythroblast stage. Middle, green: Peaks of chromatin accessibility, determined by ATAC-seq, at the promoter and in a putative intron 4 enhancer increase during terminal erythroid differentiation. Bottom, blue. Peaks of histone marks and GATA1 and TAL1 transcription factor occupancy, determined by ChIP-seq, in HSPCa and proerythroblasts. (B) Comparison of canonical *TNS1* gene to *eTNS1* short isoform (not to scale), including the unique *eTNS1* start exon (1E) highlighted in magenta. Pre-designed TaqMan primer-probe pairs (I-V) were chosen to span the canonical *TNS1* cDNA at the specific exon sites shown (Key: TaqMan Exon Probes). sgRNA (yellow triangle) location designed for CRISPR-Cas9 knockout experiments. Schematic created with BioRender.com. (C) Quantification of mRNA expression (RT-qPCR) calculated as 2(-ΔΔC(T)) for all 5 probes on days 7, 11, and 14 of erythroid cultures. Fold change in gene expression normalized to average ΔCt value for day 7 cells for each of the 5 probes, using α-tubulin as a housekeeping gene. Values are mean ± SD from 3 individual erythroid cultures. *\*p<0.05; **p<0.01; ***p<0.001; ****p<0.0001*. (D) Representative western blot for eTNS1, HA, total actin and total protein in MDCK cells (full length TNS1), CD34+ day 14 cells (eTNS1), untransfected HEK293T cells and HEK293T cells transfected with an eTNS1-HA plasmid.

Erythroblasts from day 14 cultures comprise a heterogeneous population of cells, including non-polarized, polarized, and enucleating cells. Therefore, to directly evaluate TNS1 expression in erythroblasts prior to and during polarization and enucleation, we performed immunofluorescence staining and Zeiss AiryScan confocal microscopy to examine individual erythroblasts at different stages of polarization and enucleation. TNS1 staining appeared weak and diffuse in non-polarized cells in which the nucleus had not condensed and remained centrally located (Figure 2D). In contrast, cells with a condensed nucleus polarized to one side of the cell showed abundant bright TNS1 puncta in the cytoplasm, increasing in intensity in enucleating cells (Figure 2D). Surprisingly, immunostaining for TNS1 appeared throughout the cytoplasm and was not enriched with the F-actin in the enucleosome at the rear of the nucleus in enucleating cells (Figure 2D). After nuclear expulsion, bright TNS1 puncta remained in the nascent reticulocyte, with very little TNS1 detected within the pyrenocyte (Figure 2D). This suggests that TNS1 protein levels are specifically increased late in terminal erythroid differentiation, with high expression in both polarized and enucleating cells.

### A truncated TNS1 transcript coding for a novel short isoform of TNS1 (eTNS1) is expressed in human primary erythroid cells

To investigate the origin of the short TNS1 isoform detected in human primary erythroid cells, we analyzed RNA-seq data sets from stages of terminal erythroid differentiation with respect to the genomic structure of human TNS1. The canonical human TNS1 mRNA transcript (Ensembl, *TNS1-201)* contains 33 exons encompassing 10,331 bp encoding a predicted protein of 1,735 residues with a predicted molecular weight of 185 kDa (Figures 3A, 3B, 4A). In human erythroid cells, the canonical transcript is absent. Instead, a truncated mRNA transcript of 4,292 bp encoded by 15 exons (*TNS1-208*) initiates from a novel transcription initiation site located in a novel exon 1E containing an in-frame initiator methionine, located in intron 17 of the canonical transcript (Figure 3A, 3B). This transcript encodes a predicted protein of 846 residues with a predicted molecular weight of 89 kDa. During human erythropoiesis, the truncated erythroid transcript is not expressed until early basophilic erythroblasts, increasing through terminal erythroid differentiation to very high levels in polychromatic and orthochromatic erythroblasts (Figure 3A). The area upstream of *TNS1* erythroid exon 1E has features that strongly suggest this region contains an active erythroid promoter with H3K4 trimethylation, H3K27 acetylation, and GATA1 and TAL1 occupancy (Figure 3A, Appendix Figure S5A). The features of this promoter suggest increasing chromatin accessibility during terminal erythroid differentiation paralleling increasing gene expression. In addition, an erythroid enhancer appears to be present in intron 4 of the truncated transcript, indicated by an ATAC peak, enrichment of histone H3K27 acetylation, and occupancy by the erythroid transcription factors, GATA1 and TAL1 (Figure 3A) (Xiang et al., 2024).

This truncated *TNS1-208* erythroid transcript is not present in mice or other rodents, chickens, and zebrafish (Appendix Figure S5B). Analysis of species conservation of the DNA sequence at the *TNS1-208* locus in the region of the predicted erythroid promoter, reveals that the potential GATA1 and TAL1 occupancy sites, the predicted initiator methionine ATG and the 5’ donor splice site of exon 1E, are not present in many species (Appendix Figure S5A, S6). Further, this region of the *TNS1-208* gene DNA sequence is contained within a long interspersed nuclear element (LINE). These data indicate the truncated transcript found in human erythroid cells may be primate specific.

Comparison of the canonical *TNS1* gene structure to the *eTNS1* short isoform reveal differences in addition to the unique *eTNS1* start exon 1E. Exons 20 and 21 in the canonical *TNS1* gene are missing in *eTNS1* mRNA, and the 3’UTR region of *eTNS1* is shorter when compared to canonical *TNS1* (Figure 3B). To confirm the expression of a truncated *TNS1-208* transcript in differentiated human erythroblasts, five pre-designed TaqMan probes were selected for coding exons from the 5’ to 3’ end of full-length *TNS1-201,* two of which also span exons of *TNS1-208* (Figure 3B). Probe I corresponded with the 5’ prime region, probes II, III, and IV correspond with the central region, and probe V corresponded to the 3’ prime region of *TNS1-201* (Figure 3B). Compared to day 7 cells from human erythroid cultures, only probes IV and V, which span exons in both *TNS1-201* and *TNS1-208*, showed significantly increased mRNA levels at day 14 (Figure 3C), while probes I-III showed considerably less expression. The increased *TNS1* mRNA levels detected only by probes IV and V, but not by probes I-III, strongly indicate the expression of the *TNS1-208* gene in human erythroid cells.

To determine if the protein translated from the truncated *TNS1-208* transcript shares the same molecular weight as the erythroid TNS1 protein (hereby designated eTNS1), we transfected HEK 293T cells, which lack endogenous TNS1 protein, with a recombinant human *TNS1-208* plasmid expressing eTNS1 tagged with HA at the N-terminus (Sino Biological). Western blot analysis of cell lysates from HA-eTNS1 transfected HEK 293T cells showed a prominent band at ∼125 kDa, comigrating with the CD34+ eTNS1 band (Figure 3D). These protein and mRNA expression results along with the RNA-seq analysis, indicate that human erythroid cells contain a novel short TNS1 isoform, transcribed from an alternative promoter and internal start site.

The eTNS1 protein is predicted to have 846 amino acids (Figure 4A) and a molecular weight of 89 kDa, considerably less than the apparent molecular weight of ∼125 kDa observed on SDS gels (Figure 2B). Canonical TNS1 also has an apparent molecular weight (∼220 kDa) that is greater than its predicted molecular weight (185 kDa)(Chen et al., 2000). This discrepancy is due to the low electrophoretic mobility of residues 306-981 in the TNS1 unstructured domain on SDS/PAGE (Chen et al., 2000). The presence of a portion of this unstructured central region in the eTNS1 (Figure 4A), likely contributes to the discrepancy between the predicted molecular weight of 89 kDa for eTNS1 and the apparent molecular weight of the immunoreactive band at ∼125 kDa in human erythroid cells.

**Figure 4:**
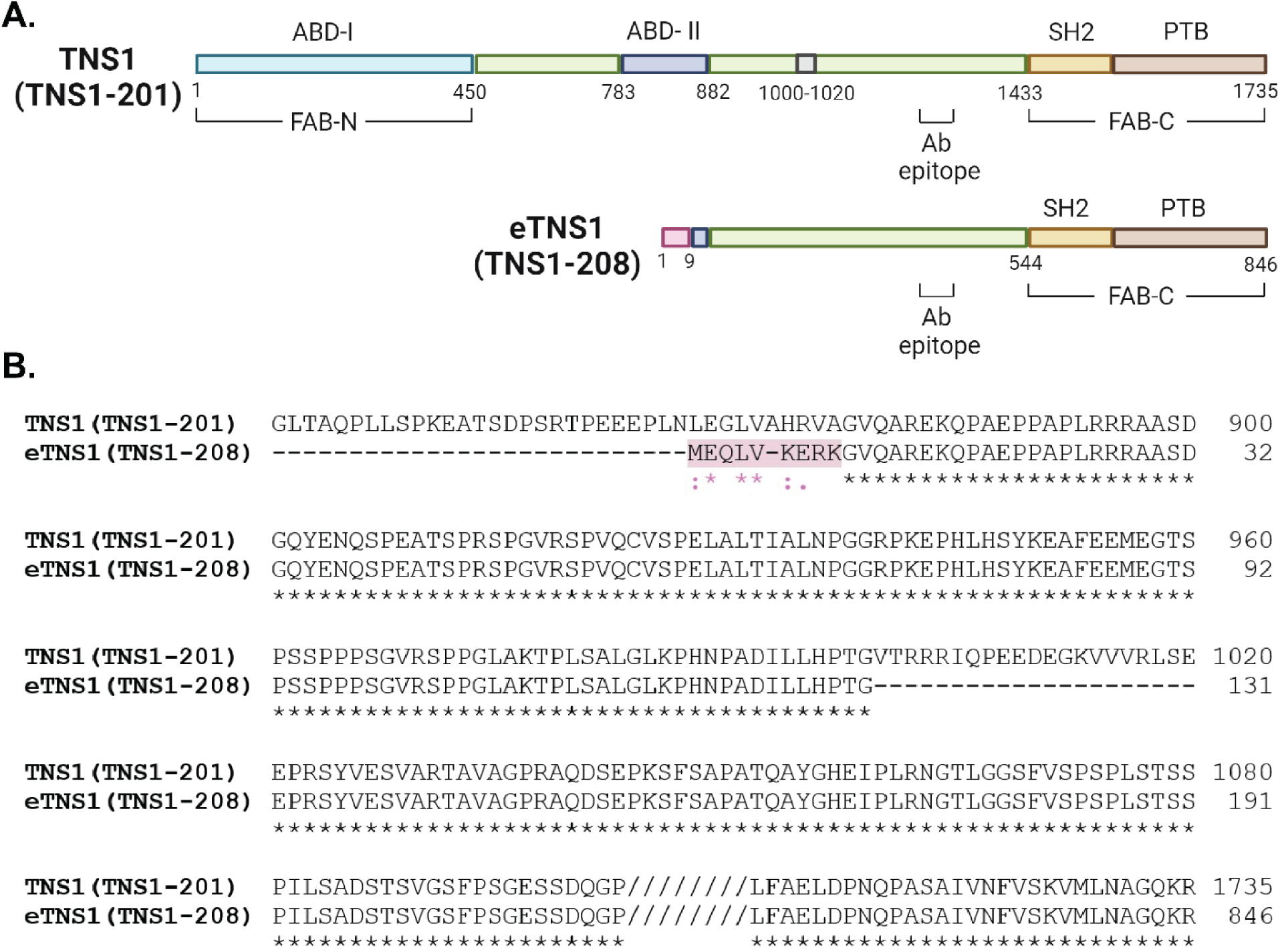
Comparison of protein domains and amino acid alignment of canonical TNS1 and eTNS1. (A) Comparison of canonical TNS1 (TNS1-201) protein domains to eTNS1 (TNS1-208). The amino acids in pink are unique to eTNS1, while the 21 amino acids in gray (1000-1020) in canonical TNS1 are missing in eTNS1. The primary antibody epitope is indicated (amino acids 1326-1339 in TNS1). Schematic created with BioRender.com. (B) Amino acid alignment using UniProt align tool showing amino acids coded for by exon 1E in eTNS1 protein, highlighted in pink. Asterisks, identical residues; Colon, residues with strongly similar properties (scoring > 0.5 in Gonnet PAM 250 matrix); Period, residues with weakly similar properties (scoring < 0.5 in Gonnet PAM 250 matrix).

Comparison of eTNS1 to full-length TNS1 protein sequences using UniProt reveals that eTNS1 is missing several N-terminal domains, including the N-terminal actin-binding domain (ABD-I) and focal adhesion targeting domain (FAB-N) (Figure 4A) (Liao & Lo, 2021). The internal transcription start site in eTNS1 results in an alternative first exon (exon 1E) coding for 9 unique amino acids that diverge from the full-length TNS1 sequence (Figure 4A,B); this alternative exon also eliminates most of the region proposed to contain an internal actin-binding domain (ABD-II) in full-length TNS1 (Liao & Lo, 2021). Following this alternative exon 1E, the eTNS1 amino acid sequence corresponds in large part to that of full-length TNS1, with an unstructured domain followed by a C-terminal focal-adhesion-binding domain (FAB-C) containing the SH2 and PTB domains (Figure 4A). Sequence alignment also revealed an additional difference in the unstructured domain of eTNS1, which is missing a 20 amino acid region present in the unstructured domain of the canonical TNS1 (Figure 4A, B). From this analysis, we can infer that terminally differentiated human erythroid cells express a unique isoform of eTNS1 transcribed from an internal transcription start site that results in a shorter protein missing the N-terminal focal adhesion-binding (FAB-N) and actin-binding domain (ABD-I), as well as the internal actin binding domain (ABD-II).

### CRISPR/Cas9 knockout of *eTNS1* in CD34+ cells reduces eTNS1 protein levels with no effect on assembly of the spectrin membrane skeleton

To investigate eTNS1 function in terminal erythroid differentiation, we introduced CRISPR/Cas9 RNP complexes into CD34+ cells on day 3 of in vitro culture (Montenont et al., 2021). We designed a guide crRNA targeting exon 24 in *TNS1-201*, which coincides with exon 6 in *TNS1-208* (Figure 3B). Genetic knockouts were confirmed by Sanger sequencing of genomic DNA on day 7 of culture using primers to amplify the region after the predicted DNA cleavage site. Although eTNS1 knockout experiments resulted in a relatively low percentage of insertions and deletions by Sanger sequencing, averaging 29% (over 7 experiments), eTNS1 protein levels were reduced by 48% on average by day 14 of culture, compared to non-transfected cells (Figure 5A, B). Western blot analysis showed that eTNS1 knockout did not result in any significant changes to total actin levels or to erythroid membrane skeleton or cytoskeletal proteins, including α1β1-spectrin, protein 4.1R and α-tubulin during terminal differentiation (Figure 5A, B). To rule out potential off-target effects due to electroporation and/or crRNA expression, we introduced CRISPR/Cas9 RNP complexes containing a non-target crRNA. Cells transfected with non-target crRNA showed no reduction in eTNS1, actin, α-tubulin or α1β1-spectrin protein levels throughout terminal differentiation, based on Western blotting (Appendix Figure S7A, B).

**Figure 5:**
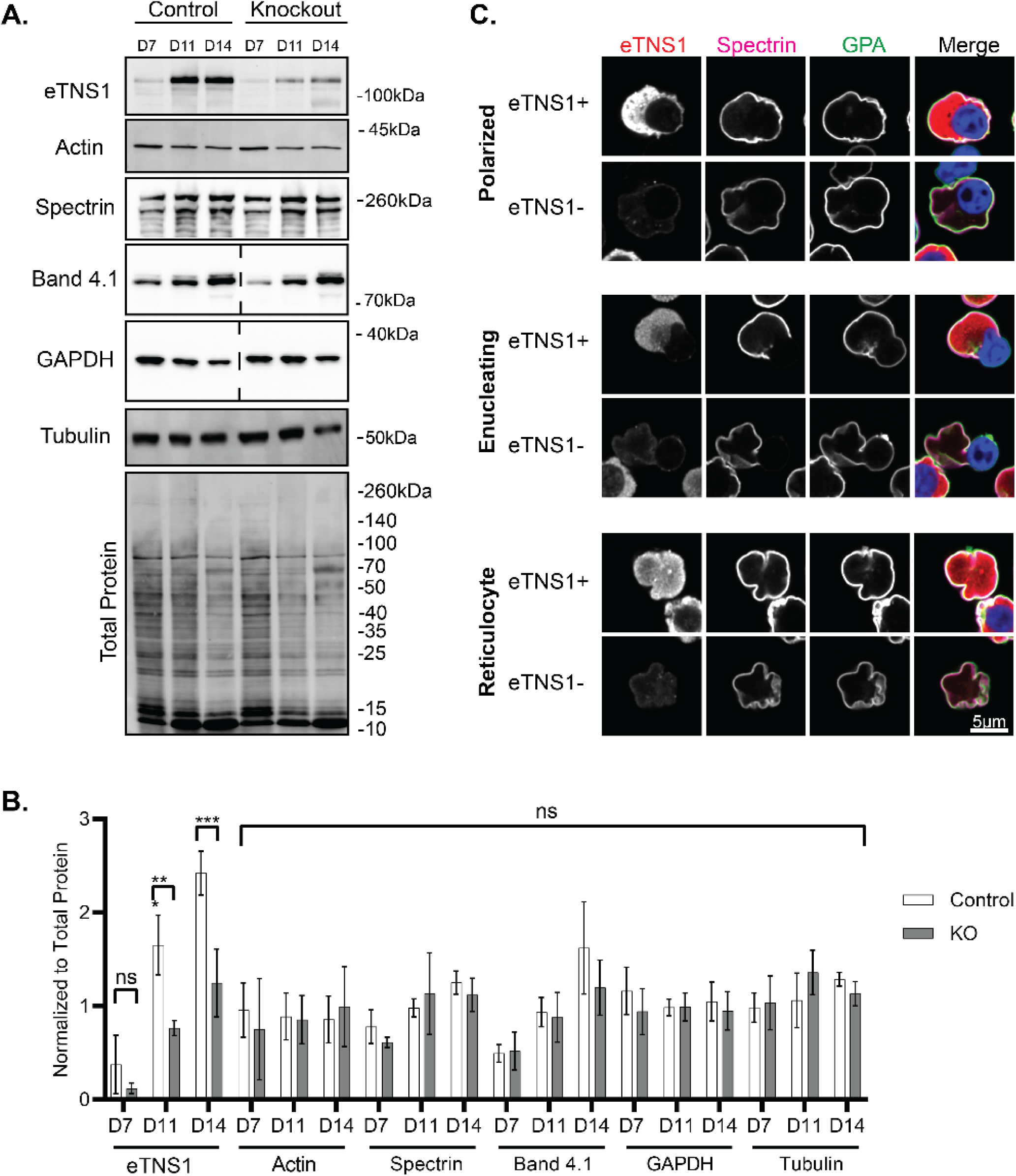
CRISPR/Cas9 knockout of eTNS1 in human erythroblasts does not affect assembly of the membrane skeleton or expression of cytoskeletal proteins. (A) Representative Western blots of eTNS1, total actin, (α1,ꞵ1)-spectrin, α-tubulin, GAPDH, protein 4.1R, and total protein from non-transfected control and eTNS1 knockout erythroid cultures. 15 µg protein loaded per lane (dashed lines indicate blots where lanes were cropped out). (B) Quantification of immunoblots normalized to total protein. Values are mean ± SD from 4 non- transfected controls and 4 knockout cultures. *\*\*p<0.01; ***p<0.001*. (C) Single optical sections of Airyscan Z-stacks of human erythroblasts from day 14 eTNS1 knockout cultures, depicting stages of enucleation, immunostained for eTNS1 (red), ꞵ1-spectrin (magenta), GPA (green), and nuclei (Hoechst; blue). Scale bar, 5µm.

We next used high resolution Zeiss Airyscan confocal microscopy to visualize eTNS1- cells and eTNS1+ cells within the same heterogeneous culture. The presence of eTNS1+ cells provided a crucial internal control to compare the morphologies side by side for cells without eTNS1 to cells with eTNS1. Airyscan confocal microscopy of eTNS1 knockout cells indicated that α1β1-spectrin assembled normally on the membrane of polarized cells, enucleating cells, and reticulocytes expressing eTNS1 (eTNS1+), or lacking eTNS1 (eTNS1-) (Figure 5C). We also noticed that some polarized and enucleating eTNS1- erythroblasts in these cultures appeared to be “stuck” in the process of nuclear expulsion, with the incipient reticulocyte assuming a dish-like or biconcave shape with a rim and a dimple, somewhat resembling that of a mature red blood cell. Single optical sections of Z-stacks as well as orthogonal XZ views of these cells show that the incipient reticulocyte of the eTNS1- cell has a distinct outer rim, while the nucleus is retained within the cell (Appendix Figure S8). By contrast, incipient reticulocytes of eTNS1+ erythroblasts do not appear to assume a biconcave-like shape while still containing their nucleus. Together with the normal expression and assembly of spectrin at the membrane (Figure 5A-C), it appeared that these eTNS1- nucleated cells were able to assemble an erythrocyte membrane skeleton that allowed the incipient reticulocyte portion of the cell to form a biconcave-like shape, without having fully completed nuclear expulsion.

### eTNS1 is required for efficient erythroblast enucleation

To determine whether loss of eTNS1 affected erythroblast enucleation efficiency, we employed single-cell microscopy analyses using the Zeiss CellDiscoverer7 (CD7) for high-throughput screening of heterogeneous cell populations. This was necessary due to the low percentage of insertions/deletions in CRISPR/Cas9 experiments. In vitro experiments were extended to day 17 to obtain cultures with a higher percentage of enucleating cells. We created a classification program to categorize cells into four distinct classes based on presence of nuclei, GPA staining, and eTNS1 staining: eTNS1+ erythroblasts, eTNS1- erythroblasts, eTNS1+ reticulocytes, and eTNS1- reticulocytes (Figure 6A-C). This analysis demonstrated that GPA intensity levels did not change in the absence of eTNS1 in differentiating erythroblasts at day 14 or day 17 (Figure 6D), suggesting that terminal erythroid differentiation is unaffected, in agreement with confocal microscopy and Western blots of membrane skeleton proteins (Figure 5A-C). Next, we investigated whether absence of eTNS1 affected the ability of erythroblasts to enucleate. Using the same classification scheme, we calculated the percentage of reticulocytes with respect to the total number of GPA-positive erythroblasts and reticulocytes. We found that the percentage of reticulocytes was significantly lower in the population of eTNS1- cells compared to eTNS1+ cells at both day 14 and day 17 in the knockout cultures (Figure 6E). This same trend was observed when comparing eTNS1- cells from knockout cultures to non-target and non-transfected control cultures (Figure 6E), leading us to conclude that cells lacking eTNS1 have an enucleation defect that impairs nuclear expulsion.

**Figure 6:**
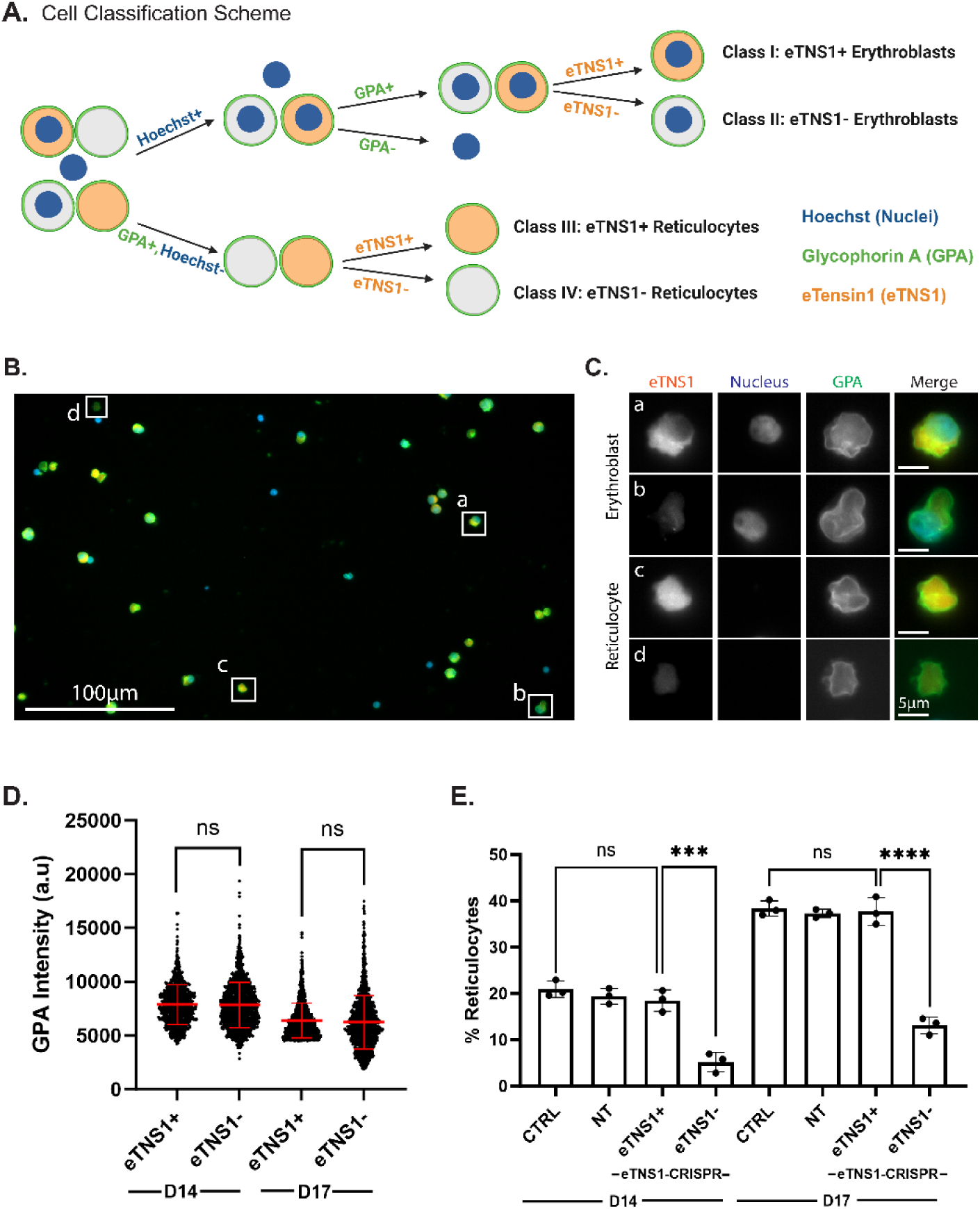
Single cell analysis reveals loss of eTNS1 impairs erythroblast enucleation. (A) Schematic of erythroblast classification from eTNS1 knockout erythroid cultures into four categories based on Hoechst (Nuclei; blue), GPA (green), and eTNS1 (orange) staining. Cells were classified into eTNS1+ erythroblasts, eTNS1- erythroblasts, eTNS1+ reticulocytes, and eTNS1- reticulocytes. Schematic created with BioRender.com. (B) Representative single tiled image showing the heterogeneous cell population from an eTNS1 CRISPR KO culture using the Zeiss CellDiscoverer 7 (CD7). Scale bar, 100 µm. (C) Representative images of individual cells depicting the four classes: (a) eTNS1+ erythroblast, (b) eTNS1- erythroblast, (c) eTNS1+ reticulocyte, and (d) eTNS1- reticulocyte. Scale bar, 5 µm. (D) Quantification of GPA intensity levels of eTNS1+ and eTNS1- cells from day 14 and day 17 eTNS1 knockout cultures. 3-5 tiled images were taken from a single coverslip for each culture. Plot reflects the mean ± SD of 1500 eTNS1+ and eTNS1- erythroblasts from three independent eTNS1 knockout cultures. (E) Quantification of eTNS1- and eTNS1+ reticulocytes in control, non-target and knockout day 14 and day 17 cultures. % reticulocytes: (# of reticulocytes) / (# of reticulocytes + # of erythroblasts). Plot reflects mean ± SD of % reticulocytes calculated from ∼6000 cells for each experiment. Three independent experiments performed for each condition. *\*\*\* p=0.0002*; *\*\*\*\* p<0.0001*.

### eTNS1 is required for F-actin assembly into the enucleosome

To further investigate the enucleation defect in eTNS1- erythroblasts, we utilized Airyscan confocal microscopy to examine F-actin organization in eTNS1 knockout erythroblasts. Maximum intensity projections of Airyscan Z-stacks showed the F-actin structures in polarized and enucleating eTNS1- cells to be extremely abnormal. Instead of F-actin accumulation into a bright focus at the rear of the nucleus (Figures 1A, 2D, 7A), eTNS1- erythroblasts had numerous mis- localized F-actin foci located in the cytoplasm or nucleus, irregular F-actin cables extending across the cell, or very little F-actin at all (Figure 7A). Quantification of F-actin intensity in eTNS1+ and eTNS1- polarized and enucleating cells using the Zeiss CD7 confirmed that eTNS1- erythroblasts in both day 14 and day 17 knockout cultures have significantly less F-actin than eTNS1+ erythroblasts (Figure 7B). This trend was maintained when F-actin intensity was normalized to GPA intensity (Figure 7C). Together, these single cell microscopy analyses suggest eTNS1 is required for the assembly of the F-actin-rich enucleosome, facilitating efficient enucleation during erythroid terminal differentiation.

**Figure 7:**
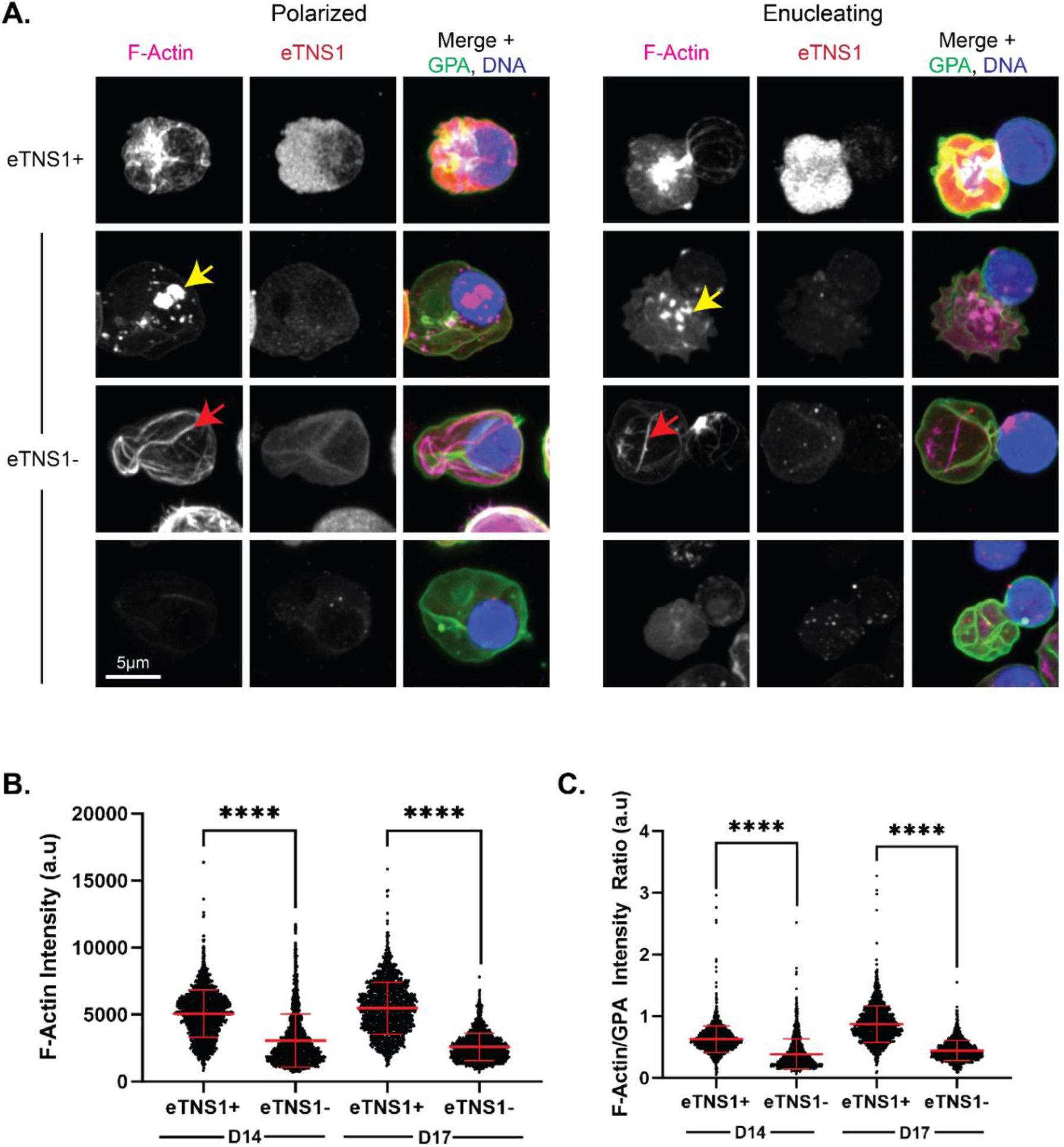
eTNS1 is required to form the F-actin enucleosome during enucleation of human erythroblasts. (A) Maximum intensity projections of Airyscan Z-stacks of polarized and enucleating human erythroblasts from day 14 cultures immunostained for F-actin (phalloidin; magenta), eTNS1 (red), GPA (green), and nuclei (Hoechst; blue). F-actin in eTNS1- cells assembled into mis-localized foci (yellow arrow), cables (red arrow), or was absent. Scale bar, 5 µm. (B) Quantification of F-actin intensity in eTNS1- and eTNS1+ erythroblasts from eTNS1 knockout cultures at days 14 and 17 using the Zeiss CellDiscoverer 7. (C) Quantification of F-actin intensity normalized to GPA intensity of eTNS1+ and eTNS1- erythroblasts from eTNS1 knockout cultures at days 14 and 17. (B-C) Plots reflect the mean ± SD of 1500 eTNS1+ and eTNS1- erythroblasts from three independent eTNS1 CRISPR KO experiments. \*\*\**\* p<0.0001*.

## Discussion

In this study, we adopted a data mining approach to identify only one ABP, TNS1, that is selectively upregulated both transcriptionally and translationally in human erythroblasts during late stages of terminal erythroid differentiation (Figure 1C, D), and confirmed this finding experimentally in human erythroid cells differentiated from CD34+ cells (Figure 2A-C). We show that the erythroid TNS1 (eTNS1) is a novel short isoform (Mr ∼125 kDa) derived from an internal transcription start site in an alternative exon 1E located in intron 17 of the canonical *TNS1* transcript (Figure 3A). Notably, we observed key differences in the gene structure of *eTNS1* (*TNS1-208*) compared to the canonical gene transcript for *TNS1* (*TNS1-201*), leading to several differences in the protein structure (Figure 4). The resulting protein is a truncated TNS1 isoform missing the FAB-N site, including the ABD-I region and PTP and C2 domains, as well as most of the ABD-II within the unstructured region (Figure 4). The remainder of the unstructured domain is present in eTNS1 (with exception of the missing 21 amino acids), as is the FAB-C site containing the SH2 and PTB domains (Figure 4). Using confocal microscopy and single-cell analyses, we demonstrated that eTNS1 is required for effective enucleation (Figure 6D) and F-actin assembly into the enucleosome (Figure 7A) but not for assembly of the spectrin membrane skeleton (Figure 5).

The *eTNS1* transcript appears to be primate specific (Figure S5A, S6A), likely due to the presence of exon 1E within a LINE leading to retrotransposition within this region of the genome (Kazazian & Moran, 2017), and is not present in mice or other rodents (Figure S5B, S6A). Since eTNS1 is required for F-actin assembly into the enucleosome in human erythroblasts (Figure 7), but mice do not express eTNS1, other actin-binding proteins may promote F-actin assembly into the enucleosome in mouse erythroblasts, such as Tmod1 (Nowak et al., 2017). Mice may also utilize additional actin-binding proteins to promote F-actin assembly into the contractile F-actin ring at the nuclear and cellular constriction, which is not present in human erythroblasts during nuclear extrusion (Figure 2C) (Newton et al., 2024; Nowak et al., 2017) (Figure 7). While the processes of terminal erythroid differentiation culminating in enucleation are broadly similar between mouse and human erythroblasts, global transcriptome analyses of human and murine erythropoiesis has revealed significant differences which may reflect divergent physiological demands and provide insights into human hematological disorders (An et al., 2015). It is intriguing to consider that loss of eTNS1 function may account for some as yet unidentified causes of human congenital dyserythropoietic anemias in which nuclear extrusion is impaired or abnormal (Iolascon et al., 2020; King et al., 2022; Risinger et al., 2019).

Nuclear extrusion is the final step in forming the nascent reticulocyte and depends on F-actin assembly and actomyosin contractility (Konstantinidis et al., 2012; Koury et al., 1989; Ubukawa et al., 2012; Wang et al., 2012). In human erythroblasts, the ABP Tmod1 plays an important role in regulating F-actin assembly and co-localizes with F-actin and NMIIB in the enucleosome (Nowak et al., 2017). Unlike Tmod1, we did not observe enrichment of eTNS1 in the enucleosome in polarized or enucleating eTNS1+ cells, or colocalization between eTNS1 and F-actin (Figure 7A), implying that eTNS1 is not a component of the enucleosome at the rear of the translocating nucleus. However, in absence of eTNS1, enucleation of erythroblasts was greatly impaired, and F-actin did not assemble into the enucleosome, but was mislocalized or not detectable (Figure 7). Since eTNS1 is missing both ABD-I and ABD-II (Figure 4) that interact with F-actin in canonical TNS1 (Lo et al., 1994), eTNS1 may have an as yet unidentified ABD that can nucleate and/or stabilize F-actin, or eTNS1 may regulate actin polymerization by activating signaling cascades mediated by SH2 and PTB domains in the conserved C-terminal domain of eTNS1.

Both the SH2 and PTB domains in canonical TNS1 bind to DLC-1, a Rho GTPase-activating protein which reduces RhoA-GTP levels, and silencing of TNS1 leads to inactivation of RhoA in some cell types (Blangy, 2017; Liao & Lo, 2021; Liao et al., 2007; Shih et al., 2015; Wang et al., 2022). Rho GTPases are critical regulators of the actin cytoskeleton and have demonstrated roles in erythroblast enucleation through multiple pathways, including Rac1/2 and CDC42 (Kim et al., 2009; Konstantinidis et al., 2012; Ubukawa et al., 2020; Ubukawa et al., 2012). It is possible that increased expression of eTNS1 and binding to DLC-1 in late stage erythroblasts may regulate actin polymerization and enucleation via activation of CDC42 (Kim et al., 2009; Ubukawa et al., 2020), or via another Rho GTPase. In addition to F-actin assembly, eTNS1 signaling may also regulate other molecular mechanisms involved in enucleation, including actomyosin contractility via NMIIB (Ubukawa et al., 2012; Wang et al., 2012; Wölwer et al., 2016) and ROCK1/MLCK (Fang et al., 2022; Konstantinidis et al., 2012; Ubukawa et al., 2012).

We demonstrate that eTNS1 is required for efficient nuclear extrusion; however, we cannot rule out that eTNS1 may also function in the terminal erythroid differentiation pathway prior to this event (Newton et al., 2024). Moreover, many of the functions/roles of eTNS1 postulated here are based on the structural similarity to TNS1. However, despite high sequence conservation of eTNS1 with the C-terminal portion of TNS1, eTNS1 may have unique functions due to spatiotemporal localization and unique binding partners in erythroid cells, as seen with other members of the TNS family in different cell types (TNS1-4) (Blangy, 2017; Liao & Lo, 2021). Future studies investigating the binding partners and function of the conserved domains in eTNS1 are required to explore these ideas further.

In summary, our study identifies a novel isoform of TNS1 (eTNS1) expressed in late stages of human terminal erythroid differentiation that localizes to the cytoplasm during nuclear polarization and enucleation. We demonstrate that eTNS1 is required for F-actin polymerization into the enucleosome, and for efficient enucleation of human erythroblasts to generate reticulocytes. Elucidating the role of eTNS1 in the molecular pathways underlying this complex cellular process will provide insights into as yet unappreciated species-specific functional differences between mouse and human erythroid cell physiology, and may aid in improving human erythroid culture systems to produce erythrocytes ex vivo. Future studies will aim to further characterize the function of eTNS1, its potential binding partners, and how eTNS1 regulates the actin cytoskeleton during human erythroid terminal differentiation and enucleation.

## Methods

### Data mining strategy to identify actin-binding proteins up-regulated in late stages of terminal erythroid differentiation

To understand the role of F-actin in erythroblast enucleation, we first curated a list of 135 ABPs and NFs across 20 actin-associated protein families (Campellone & Welch, 2010; Chesarone & Goode, 2009; Fowler, 2013; Siton-Mendelson & Bernheim-Groswasser, 2017), as well as a control group of 12 known red cell membrane skeleton proteins and 25 red cell transmembrane and membrane-associated proteins (Fowler, 2013; Lux, 2016) (Appendix Table S1). These lists were utilized in our data mining strategy to cross reference two well-established RNA-seq (Yan et al., 2018) and proteomics (Gautier et al., 2016) datasets for proerythroblast, basophilic erythroblast, and orthochromatic erythroblast stages of human erythroblast differentiation derived from CD34+ cells. Hits from the cross-referencing method were visualized by plotting fold-change, normalized to the proerythroblast stage of differentiation on a linear scale (log 2- fold change). The threshold for increased expression was set at >1.0-fold increase, compared to proerythroblast expression levels (Appendix Table S2).

### CD34+ cell culture and erythroid differentiation

CD34+ cells were isolated from cord blood (STEMCELL Technologies, 70008.2) and differentiated along the erythroid pathway using a previously described three-phase culturing system (Dulmovits et al., 2016). Briefly, CD34+ cells were thawed in IMDM (Gibco 12440053) with 5% FBS (Genesee Scientific 25-550H) and sub-cultured on days 0, 4, 7, 11, and 14. On days 0 and 4 (phase 1), cells were seeded at 1x10^5^ cells/mL in IMDM supplemented with 3% AB serum (Fisher Scientific, BP2525100), 2% PB plasma (STEMCELL Technologies, 70039), 10 µg/mL insulin (Sigma-Aldrich, 91077C), 3 U/mL erythropoietin (Prospec, CYT-201), 3 U/mL heparin (Sigma-Aldrich, H3149), 200 ug/mL transferrin (Fisher Scientific, 50-489-920), and 10 ng/mL stem cell factor (SCF) (STEMCELL Technologies, 78062), 1 ng/mL IL-3 (STEMCELL Technologies, 78040) and 1% Penicillin-Streptomycin (Gibco 15140122). On day 7 (phase 2), cells were seeded at 1x10^5^ cell/mL in the above media formulation without IL-3 and Pen-Strep. On day 11 and 14 (phase 3), cells were seeded at 1x10^6^ cells/mL in the above media formulation without IL-3, Pen-Strep or SCF, and transferrin was increased to 1 mg/mL.

### Giemsa staining

3x10^5^ cells were removed from culture on days 7, 11, and 14 of in vitro erythroid differentiation. Cells were washed once by centrifugation at 200 g for 10 minutes, resuspended in 200 µL of PBS (Genesee Scientific, 25-507B), transferred to an EZ Single Cytofunnel (Eprediam, A78710003), and cytospun onto glass slides at 1200 rpm for 3 minutes using the Thermo Scientific Cytospin 4 centrifuge. Samples were then air dried before fixation in 100% methanol (Fisher Scientific, A4544) for 5 minutes. Samples were stained using Modified Giemsa Stain (Sigma, GS5000) diluted 1:20 in water for 45 minutes. Samples were air dried before adding 30 µL Permount^TM^ Mounting Medium (Fisher Scientific, SP15-100) followed by a coverslip. Samples were air dried overnight at room temperature before imaging. Images were acquired with a Zeiss Axio Observer 7 wide-field inverted microscope (63x oil objective, NA 1.4) using an Axiocam 208 color camera. All images were processed using Zen software (Carl Zeiss) for further analysis.

### Staining for flow cytometry

1x10^6^ cells were removed from culture on days 7, 11, 14, and 17 of in vitro erythroid differentiation and washed twice in FACS Buffer (PBS + 2% FBS) by centrifugation at 300 g for 5 minutes at 4°C. After washes, cells were resuspended in 100 µL of FACS Buffer and stained for 15 minutes at room temperature with an antibody cocktail containing anti-CD71 (1:50; PerCP/Cyanine 5.5; BioLegend, 334113), anti-CD235a (1:500; PE/Cyanine 7; BioLegend, 349111), anti-Band 3 (1:400; FITC; kindly provided by Dr. Xiuli An, New York Blood Center), anti-CD49D (1:33; PE; Miltenyi, 130-118-548), anti-CD44 (1:33; BV510; BioLegend, 103043), anti-CD47 (1:33; BV605; BD Biosciences, 563749), DRAQ5 (1:20,000; BioLegend, 424101), and SYTO^TM^ 60 Red (1:20,000; Thermo Fisher Scientific, S11342). After staining, cells were washed once before resuspending cells in 300 µL FACS Buffer. Compensation beads (BioLegend, 424602) were used along with Cell Staining Buffer (BioLegend, 420201) for single color controls per the manufacturer’s instructions. Samples were analyzed by a BD FACSAria Fusion High-Speed Cell Sorter and FCS Express 7 (Denovo Software).

### TaqMan gene expression analysis

Cells were removed from culture on days 7, 11, and 14 of in vitro erythroid differentiation and washed twice in PBS. RNA extraction was performed using GeneJET RNA Purification Kit (Thermo Fisher Scientific, K0731) according to manufacturer’s instructions. Reverse transcription was performed on equal amounts of total RNA for each sample using SuperScript^TM^ IV Reverse Transcriptase (Thermo Fisher Scientific, 18090050) to synthesize cDNA. RT-qPCR was performed using TaqMan^TM^ Fast Advanced Master Mix (Thermo Fisher Scientific, 4444556) in 20 µL reactions using equal amounts of cDNA per sample mixed with pre-designed TaqMan probes (Thermo Fisher Scientific; see Reagents and Tools Table for probe information). All TNS1 TaqMan probes were used with an FAM reporter dye, while the α-tubulin probe (housekeeping gene) was used with a VIC reporter dye. Reaction composition and cycling conditions followed the manufacturer’s instructions in a 96-well fast read plate (Genesee Scientific, 24-310) using the Azure Cielo 3 Real Time PCR System (Azure Biosystems, AIQ030).

### Transient transfection of HEK 293T cells with HA-eTNS1 plasmid

*TNS1-208* (eTNS1) was subcloned into mammalian expression vector pCMV3-N-HA (Sino Biological). Plasmid DNA was introduced into DH5ɑ competent cells (Thermo Fisher Scientific, 18258012) by heat shock method. 50 ng of plasmid was added to 50 µL competent cells and allowed to incubate on ice for 30 minutes, followed by a 45 second heat shock at 42℃, then immediately placed on ice for 5 minutes. 200 µL of SOC medium (Thermo Fisher Scientific, 15544-034) was then added and incubated while shaking at 37°C for 1 hour. Transformed cells were then spread on LB agar plates (MP Biologicals, 3002-231) containing 50 µg/mL kanamycin (Fisher Scientific, BP906-5). Plates were incubated overnight at 37°C. Single colonies were chosen the following day, transferred to 5 mL of LB broth (Fisher Scientific, BP1426-2), and incubated shaking overnight at 37℃. Plasmid DNA was isolated using a mini plasmid purification kit (Thermo Fisher Scientific, K2100-01).

HEK 293T cells (ATCC, CRL 3216) were grown to confluence in DMEM (Genesee Scientific, 25- 500) with 10% FBS (Genesee Scientific, 25-550H) and 1% Penicillin-Streptomycin (Gibco, 15140122). Cells were transfected using Lipofectamine 3000 (Thermo Fisher Scientific, L3000001) according to manufacturer’s instructions. Briefly, Lipofectamine 3000 reagent, P3000 reagent and plasmid DNA (1 µg/well of 6-well plate) were mixed in Opti-MEM (Thermo Fisher Scientific, 31985070) and allowed to incubate for 15 minutes at room temperature before being added to cells. Cells were incubated at 37°C for 6 hours at 5% CO2 before collection for Western blotting.

### Western blotting

Whole cell lysates were prepared using radioimmunoprecipitation assay buffer (150 mM NaCl, 1% Nonidet P-40, 0.5% sodium deoxycholate, 0.1% sodium dodecyl sulfate, 1 mM EDTA and 50 mM Tris HCl, pH 8.0) (Hu et al., 2013) with protease inhibitor cocktail (1:100; Thermo Fisher Scientific, 78429) and phosphatase inhibitors (1:100; Thermo Fisher Scientific, 78420). Samples were sonicated with a probe sonicator (Qsonica, Q55 Sonicator) for 10 seconds and passed through a shredder column (Lamda Biotech, PROT-SC) before determining protein concentration by BCA Protein Assay (Thermo Fisher Scientific, 23250). Cell lysates were then diluted 1:1 in 2x Laemmli sample buffer (Bio-Rad, 1610737) and boiled for 5 minutes at 95°C. 15 µg total protein per sample was loaded onto a 4-20% or 4-12% linear gradient SDS-PAGE mini-gel (Thermo Fisher Scientific, XP04200BOX, XP04120BOX) and electrophoresed at 150V for 1 hour in 1x running buffer (25 mM tris base, 192 mM glycine, 0.1% SDS), transferred to a PVDF membrane (Millipore Sigma, IPFL00010) in 1x Bjerrum semi-dry transfer buffer (48 mM Tris base, 39 mM glycine, 0.0375% SDS, 20% methanol) using a Trans-Blot Turbo Transfer System (Bio-Rad 1704150). Total protein staining was performed using the Revert 700 Total Protein Kit (LI-COR, 926-11016) and scanned with a Chemidoc MP Imaging System (Bio-Rad 12003154). Blots were then blocked with Intercept (PBS) Blocking Buffer (LI-COR, 927-70001), then incubated with primary antibodies (see Reagents and Tools Table) diluted in Intercept (PBS) Blocking Buffer overnight at 4°C with gentle rocking. The blots were then washed 3x for 5 minutes each in TBST (20 mM tris base, 137 mM sodium chloride, 0.1% tween-20, pH 7.6) and incubated with IRDye 680RD donkey anti-mouse IgG (LI-COR, 926-68072) and IRDye 800CW donkey anti-rabbit IgG (LI-COR, 925-32213) at 1:20,000 diluted in Intercept (PBS) Blocking Buffer and incubated for 2 hours at room temperature with gentle rocking. Blots were then washed again 3x for 5 minutes each before being scanned with a Chemidoc MP Imaging System (Bio-Rad 12003154). The band intensities were quantified using ImageJ and normalized to total protein staining.

### Fluorescence staining and confocal microscopy

1x10^6^ cells in 200 µL were taken out of CD34+ cultures on different days during in vitro erythroid differentiation and placed on a 12 mm fibronectin-coated coverslip (Neuvitro, GG-12-1.5) and incubated at 37°C for 2-3 hours. Media was aspirated followed by one 1x PBSi wash (200 µL PBS per coverslip) before fixation with 200 µL 4% paraformaldehyde (PFA) (Electron Microscopy, 15710) per coverslip and incubating at room temperature overnight in the dark. After fixation, cells were washed 3x with PBS, permeabilized with 0.3% Triton X-100 (Sigma-Aldrich, 11332481001) in 1x PBS for 15 minutes and blocked with 3% BSA (Genesee Scientific, 25-529) and 1% normal goat serum in PBS (blocking buffer) for 2 hours at room temperature. Rabbit anti-Tensin 1 antibody (affinity-purified polyclonal antibody (aa 1326-1339) prepared by Dr. Su Hao Lo, University of California, Davis) was used at 1:100 dilution in the blocking buffer and incubated at room temperature for 2 hours. Cells were then washed 3x in PBS, 5 minutes per wash, before adding secondary antibody solution and incubating at room temperature for 1 hour. Secondary antibody solution contained Alexa 594-conjugated goat anti-rabbit (1:400; Thermo Fisher Scientific, A-11012), Alexa 647-Phalloidin (150 nM; Thermo Fisher Scientific, A-22287), Hoechst 33342 (1:500; Thermo Fisher Scientific, H3570), and FITC-conjugated mouse anti-human GPA (1:100; BD Biosciences, 559943). Cells were then washed 3x with PBS for 5 minutes per wash, and mounted using ProLong^TM^ Glass Antifade Mountant (Thermo Fisher Scientific, P36980). Airyscan Z-stack images were acquired using a Zeiss LSM-880 laser scanning confocal microscope (63x oil objective, NA 1.4) with a 0.17µm Z-step. All images were processed using Zen blue version 2.6 (Carl Zeiss) for further analysis.

### Bioinformatic analyses

Human erythroid mRNA expression data were obtained from (An et al., 2014)(GEO GSE53983) and (Schulz et al., 2019)(GEO GSE128269). Human erythroid ATAC-seq data were obtained from (Schulz et al., 2019)(GEO GSE128269). ChIP-seq data of GATA1 and TAL1 occupancy in human erythroid cells was obtained from (Huang et al., 2016)(GEO GSE70660). TNS1 transcripts were obtained from Ensembl Genome assembly: GRCh37.p12 (Martin et al., 2023). Multi-species Multiz alignment and conservation of the TNS1 locus in 100 vertebrates was obtained from the UCSC Genome Browser database (Blanchette et al., 2004; Nassar et al., 2023).

### CRISPR/Cas9 genome editing

CD34+ cells were transfected with CRISPR/Cas9 complexes on day 3 of culture, as previously described with some modifications (Montenont et al., 2021). Equal portions (2.5 µL each) of Alt-R CRISPR-Cas9 tracrRNA 5 nmol (IDT, 1072532) and pre-designed TNS1 Alt-R CRISPR-Cas9 CRISPR RNA (crRNA; IDT) (4304-4323bp: ATCGGAGACCCACACTGTCCCGG) or Alt-R CRISPR-Cas9 negative control crRNA #1 (IDT, 1079138) for non-targeting control were mixed in a polymerase chain reaction tube and heated to 95°C in a thermal cycler, then cooled to room temperature for 10 minutes. Ribonucleoprotein complex (RNP) was then formed by mixing the following in a PCR tube: 1.2 µL tracrRNA-crRNA duplex, 2.1 µL PBS (no calcium or magnesium), and 1.7 µL Alt-R *Streptococcus pyogenes* Cas9 V3 (IDT, 1081058), heated in a thermal cycler at 37°C for 4 minutes and cooled to room temperature for 10 minutes. At the same time, cells were counted and washed 2x in sterile PBS spinning at 200 x g for 10 minutes (0.25x10^6^-4x10^6^ total cells). After the last wash, the cell pellet was resuspended in 20 µL of pre-warmed P3 primary cell solution (Lonza Kit, V4XP-3032), 2.5 µL RNP complex, and 1 µL Alt-R Cas9 electroporation enhancer 2 nmol (IDT, 1075915). After mixing, the cell mixture was transferred to a 16-well electroporation strip and transfected using an Amaxa 4D nucleofector program EO100 or DZ100. After electroporation, 75 µL of pre-warmed phase 1 media was added to the cells in the strip and incubated for 5 minutes at room temperature. Cells were then transferred to a 24-well plate containing 500 µL phase 1 media and incubated at 37°C, at 5% C02. 24 hours later (day 4), cells were passaged and re-plated in phase 2 media, as described above.

### DNA analysis for Sanger sequencing

5x10^5^ cells were removed from culture on day 7 for DNA analysis. After washing cells once with PBS, genomic DNA was extracted from cells by resuspending the pellet in 250 µL of QuickExtract DNA Extraction Solution (Lucigen, QE09050). Samples were heated at 65°C for 6 minutes and 98°C for 2 minutes before amplifying the region surrounding the CRISPR cut site by PCR. Q5 Hot Start High-Fidelity 2x Master Mix (New England Biolabs, M0494S) was used for PCR amplification with primers 5’-CCATGGTAGCACTGTCTCCAGC-3’ and 5’-CTGGCATGGAGTACTTGGAG-3’.

All samples were run on a 1.5% agarose gel with SYBR Safe DNA Gel Stain (Thermo Fisher Scientific, S33102) for DNA visualization. DNA was extracted from the agarose gel using Zymoclean Gel DNA Recovery Kit (Genesee Scientific, 11-300) before being sent for Sanger sequencing.

### Imaging with ZEISS CellDiscoverer 7

1x10^6^ cells were fixed in 4% PFA and immunostained as described above. Images were acquired with a Zeiss CellDiscoverer 7 confocal fluorescence microscope using the plan-Apochromat 50x/1.2 water immersion objective lens. 54 tiles of images (375.36 μm x 226.32 μm) were collected and further processed by stitching together to form one full image (2.07 mm x 1.86 mm), which was then analyzed using a custom analysis program to distinguish erythroblasts and reticulocytes. First, Hoechst staining was used to identify erythroblasts and pyrenocytes. Then, GPA+ erythroblasts were classified separately from GPA- pyrenocytes. Erythroblasts were then further classified into eTNS1+ erythroblasts and eTNS1- erythroblasts. Because reticulocytes have no nuclei, all GPA+ cells were identified initially, followed by identification of cells with no Hoechst staining, *ie.*, reticulocytes. Then, we classified the reticulocyte group into eTNS1+ reticulocytes and eTNS1- reticulocytes. The percentage of eTNS1+ and eTNS1- reticulocytes with respect to eTNS1+ and eTNS1- erythroblasts, was calculated as follows: (total number of reticulocytes) / (total number of reticulocytes + total number of erythroblasts) for either the eTNS1+ or eTNS1- categories. Thresholds were set manually for samples from knockout cultures (eTNS1- cells and eTNS1+ cells), cultures transfected with non-target crRNA, and non-transfected control cultures. To measure the F-actin intensity of eTNS1+ and eTNS1- erythroblasts, an additional parameter for rhodamine-phalloidin intensity was added to the existing program.

### Statistical analysis

Each experiment was replicated on at least three separate occasions to provide at least three biological replicates. Unpaired t-tests were used to detect differences between two groups of data. To determine differences between multiple groups of data, a one-way ANOVA analysis was performed. All statistical analysis was conducted using GraphPad Prism 10.

## Author contributions

**AG**: Conceptualization; Data curation; Formal analysis; Methodology; Visualization; Investigation; Validation; Writing-original draft; Writing-review and editing. **MC**: Data curation; Formal analysis; Investigation; Validation; Visualization; Methodology; Writing-original draft; Writing-review and editing. **DD**: Data curation; Formal analysis; Investigation; Validation; Visualization; Writing-review and editing. **SB**: Data curation; Investigation; Validation; Formal analysis. **VS**: Data curation; Formal analysis; Investigation; Visualization; Writing-review and editing. **PG**: Data curation; Formal analysis; Visualization; Writing-review and editing. **SHL**: Conceptualization; Methodology; Writing-review and editing. **VMF**: Conceptualization; Supervision; Methodology; Project administrator; Funding acquisition; Writing-review and editing.

## Acknowledgements

We thank Lio Blanc and Julien Papoin at the Feinstein Institute for Medical Research for initial assistance with human erythroid cell cultures, Jack Mason at the University of Delaware with Western blots and RT-PCR. We also thank Jaysheel D. Bhavsar and Shawn W. Polson for assistance with initial RNA-seq analysis in the Bioinformatics Data Science Core Facility at the University of Delaware’s Center for Bioinformatics and Computational Biology, supported by Delaware INBRE (NIGMS P20GM103446), NIH Shared Instrumentation Grant (NIGMS S10OD028725), the State of Delaware, and the Delaware Biotechnology Institute, which also supported the BioStore data management resource at the University of Delaware Bioinformatics Data Science Core used in this study. This research was also supported by a grant from NIH/NHLBI (R01HL083464) (VMF), funds from the University of Delaware (VMF), a Core Access Award from Delaware INBRE (NIGMS P20GM103446) (VMF) for University of Delaware Bioimaging Center and Flow Cytometry Core use, and a grant from NIH/NIDDK (R01DK111539) (PG).

---

**Figure S1:**
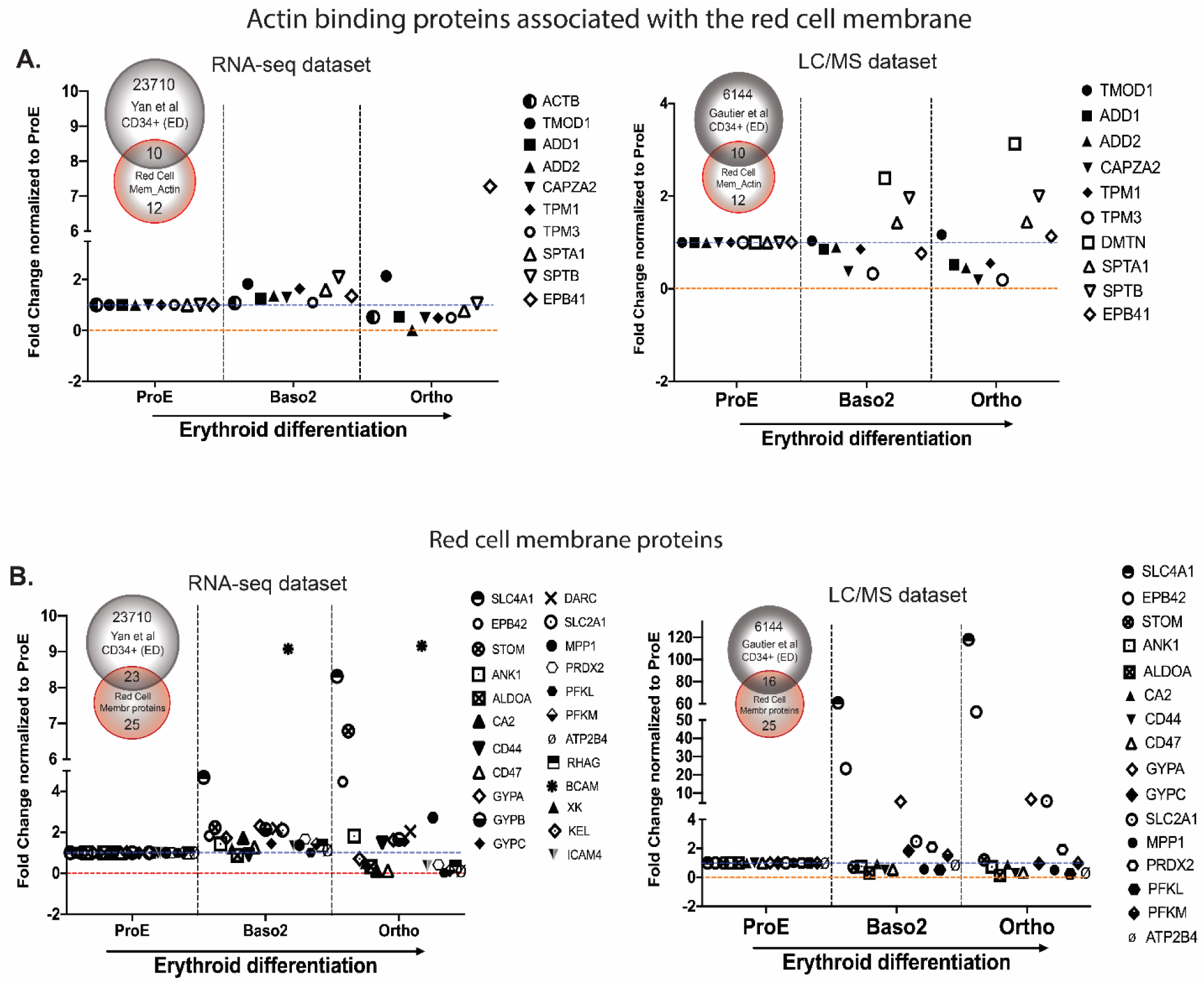
Database mining for known ABPs associated with the mature red cell membrane skeleton, and other red cell membrane proteins, as benchmarking for our search strategy for novel ABPs/NFs. Using our curated list of ABPs associated with the red cell membrane skeleton and other known red cell membrane-associated proteins (Appendix Table S1B), changes in mRNA expression levels were assessed from proerythroblast (ProE), basophilic erythroblast (Baso2), and orthochromatic erythroblast (OrthoE) stages. (A) Within the mature red cell membrane skeleton proteins dataset, most of the genes did not display an increase in mRNA expression at the OrthoE stage with respect to the ProE stage. The proteomics (LC-MS) dataset confirmed well-known expression changes for various red cell membrane skeleton proteins including tropomodulin (TMOD1), DMTN, and spectrins (SPTA1, SPTB). (B) Various known red cell transmembrane and membrane-associated proteins were found to be transcriptionally and translationally upregulated at the OrthoE stage (SLC4A1, EPB42). This demonstrated that our database mining methodology provided appropriate benchmarking for analyses of a test group of other ABPs/NFs (Figure 1).

**Figure S2:**
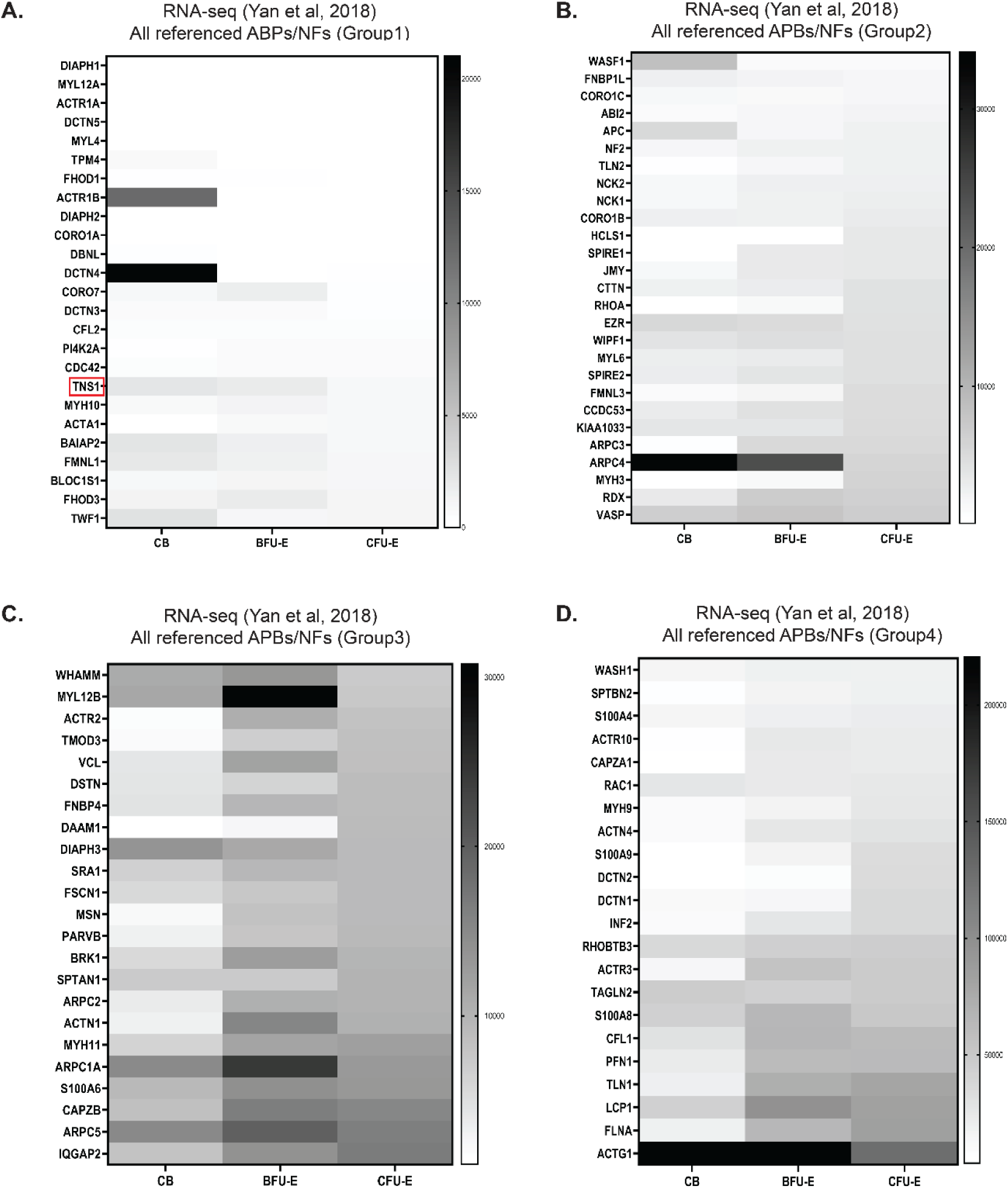
Analysis of expression levels for 97 ABP/NF hits from the RNA-seq dataset reveals *TNS1* mRNA levels remain low during early stages of erythroid differentiation. The RNA-seq dataset hits were unbiasedly clustered arbitrarily into 4 separate groups for optimal visualization via heatmap on GraphPad Prism v10 (cluster 1 = 25 genes, cluster 2 = 27 genes, cluster 3 = 23 genes, cluster 4 = 22 genes). All clusters display raw counts from early stages of erythroid differentiation (CB = Cord Blood; BFU-E = Burst-Forming Unit-Erythroid; CFU-E = Colony-Forming Unit-Erythroid), with higher expression indicated by a darker rectangle. The majority of the 97 ABP/NF hits, including TNS1 (red box), showed no significant change in mRNA expression at these early stages of erythroid differentiation.

**Figure S3:**
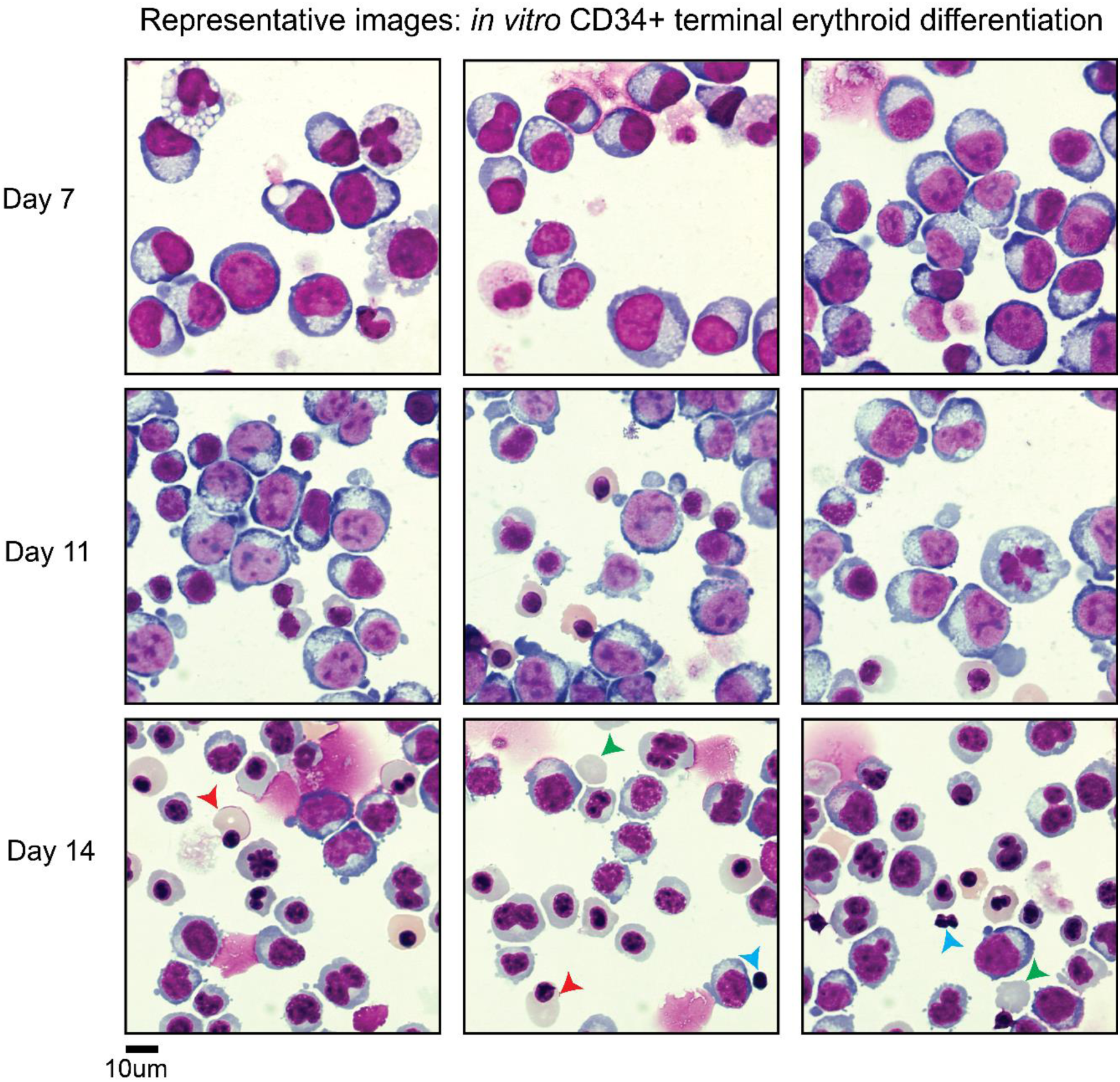
Morphological analysis confirms terminal erythroid differentiation of human CD34+ cells in vitro. Representative images of Wright-Giemsa stained CD34+ cells at days 7, 11, and 14 during in vitro erythroid differentiation show expected morphological changes of maturing erythroblasts. Scale bar, 10 µm. Day 7 cultures are primarily composed of larger cells with uncondensed nuclei. By day 11, smaller cells with condensed nuclei either centrally located or polarized to one side of the cell can be seen. Although these cultures remain heterogeneous, by day 14, cells begin to enucleate (red arrows) resulting in reticulocytes (green arrow) and pyrenocytes (blue arrow).

**Figure S4:**
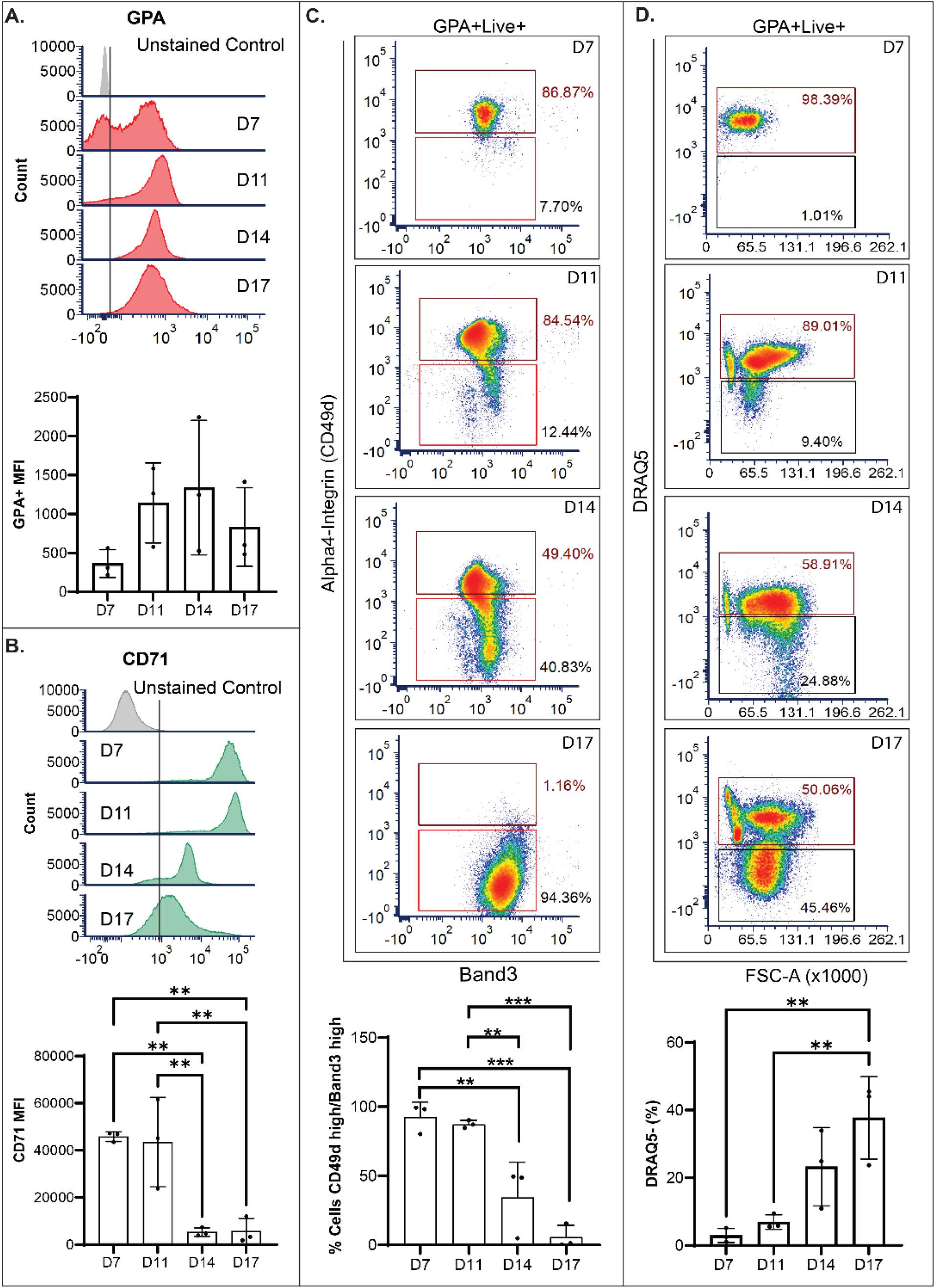
Flow cytometric analysis of human CD34+ cells in erythroid cultures confirms surface marker characteristics of terminally differentiating erythroblasts. (A-B) Histograms and quantification of mean fluorescence intensity for glycophorin A (GPA) and transferrin receptor (CD71) levels in differentiating (control) CD34+ cells. Unstained cells used as a gating control. Only live cells were analyzed. © Live GPA-positive cells analyzed for CD49d (α4- integrin) and band3 expression. Gates set for CD49d-high, band3-hi (maroon), and graphed as a change in percentage population across 3 biological replicates. (D) Live GPA-positive cells analyzed for enucleation efficiency using DRAQ5 dye, gated for DRAQ5-positive (maroon) and DRAQ5-negative (black gate) cells and graphed as a change in percentage of population for DRAQ5 negative across 3 biological replicates. Values are mean ± SD from 6 individual erythroid cultures. *\*p<0.05; **p<0.01; ***p<0.001*.

**Figure S5:**
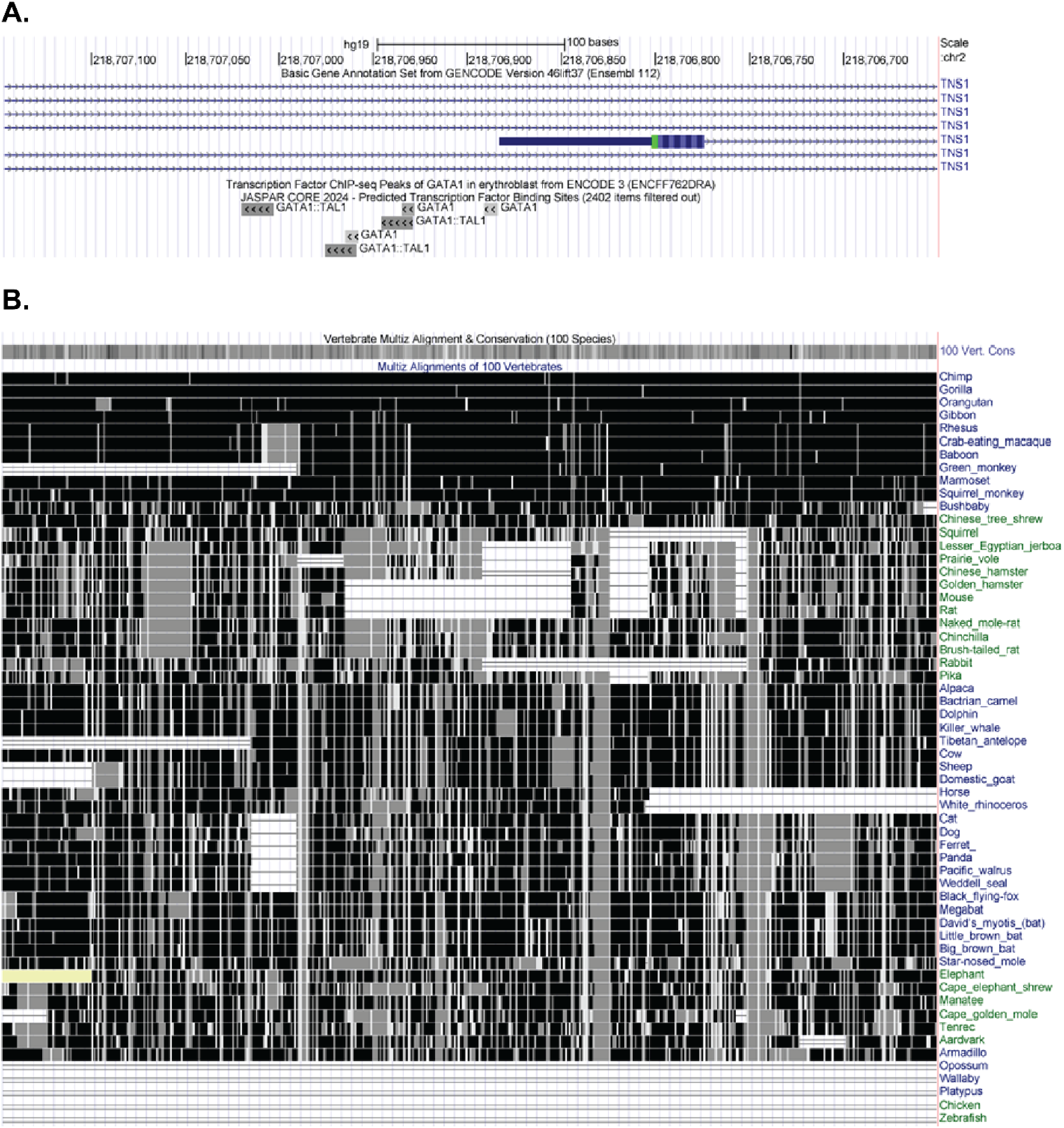
Conservation at the TNS1 locus across vertebrate species. (A) The region encoding the human erythroid exon 1E and 5’ flanking putative promoter region is shown. Exon 1E is shown as a blue bar. The location of the initiator methionine of the erythroid TNS1 peptide (eTNS1) in exon 1E is shown in green. Candidate transcription factor binding sites for GATA-1 and TAL-1 are indicated below. (B) Multi-alignment analysis of 100 vertebrates reveals strong conservation among primates. Regions deleted from genomes of individual species are shown in white.

**Figure S6:**
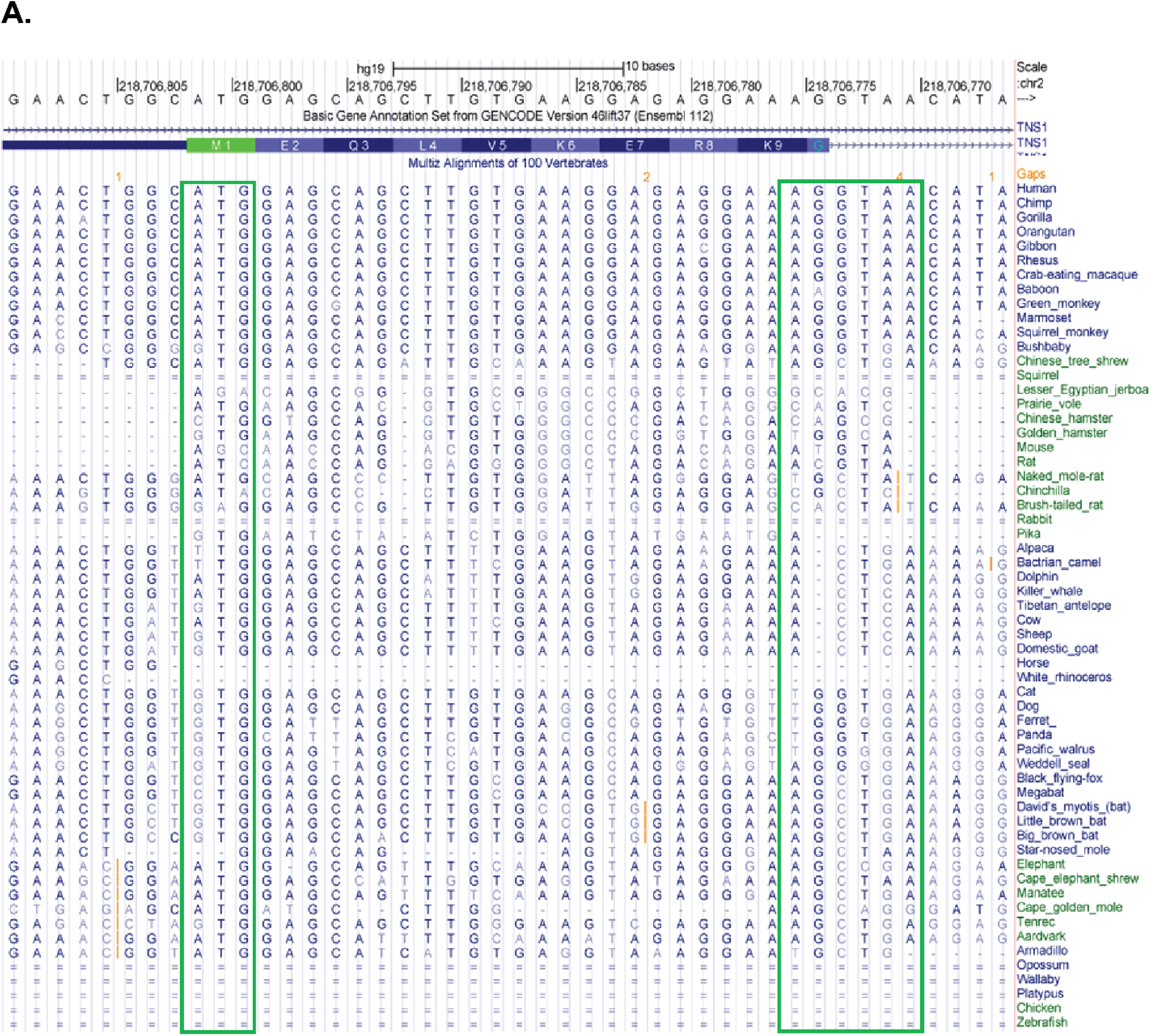
Nucleotide-level analysis at the TNS1 locus across vertebrate species. The region encoding the human erythroid exon 1E is shown. The location of the initiator methionine for the erythroid TNS1 protein (eTNS1) in exon 1E is indicated in green. There is strong conservation of the 5’ untranslated region, initiator methionine, coding region, and 5’ donor splice site among primates in multi-alignment analysis of 100 vertebrates. Other species lack the upstream 5’ untranslated region, the “A” of the initiator methionine “ATG” and/or the 5’ donor splice site consensus sequence, AG:gttaa.

**Figure S7:**
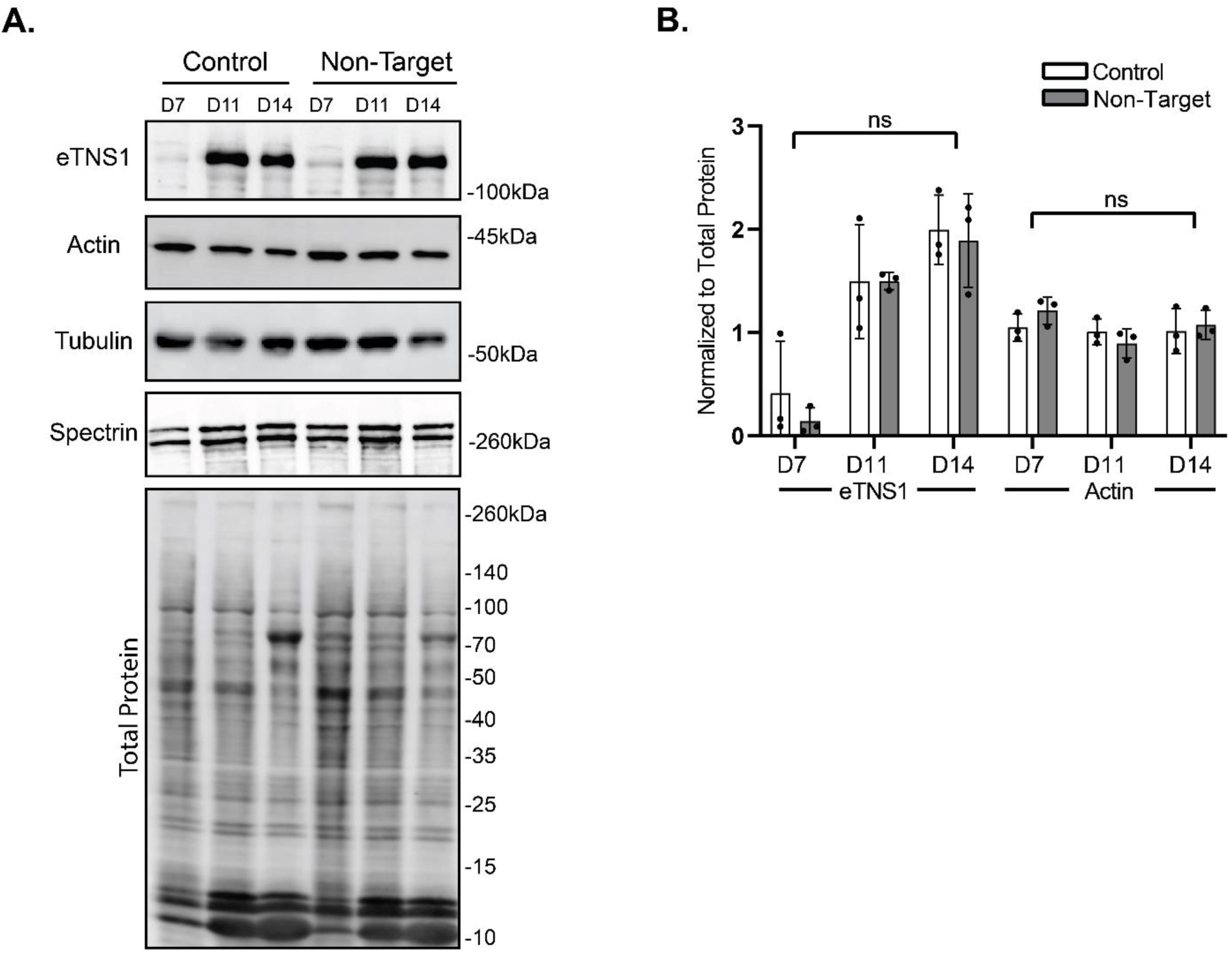
Introduction of CRISPR/Cas9 RNP complexes containing non-target crRNA had no effects on eTNS1, total actin, α-tubulin, or erythroid spectrin protein expression. (A) Representative Western blots of eTNS1, total actin, α-tubulin, (α1β1)-spectrin, and total protein during in vitro terminal erythroid differentiation of CD34+ cells in non-transfected control cultures compared to non-target cultures. 15 µg protein loaded per lane. (B) Quantification of eTNS1 and total actin in control compared to non-target cells, normalized to total protein. Band intensities were analyzed by ImageJ. Values are means ± SD from 3 individual CD34+ erythroid cultures.

**Figure S8:**
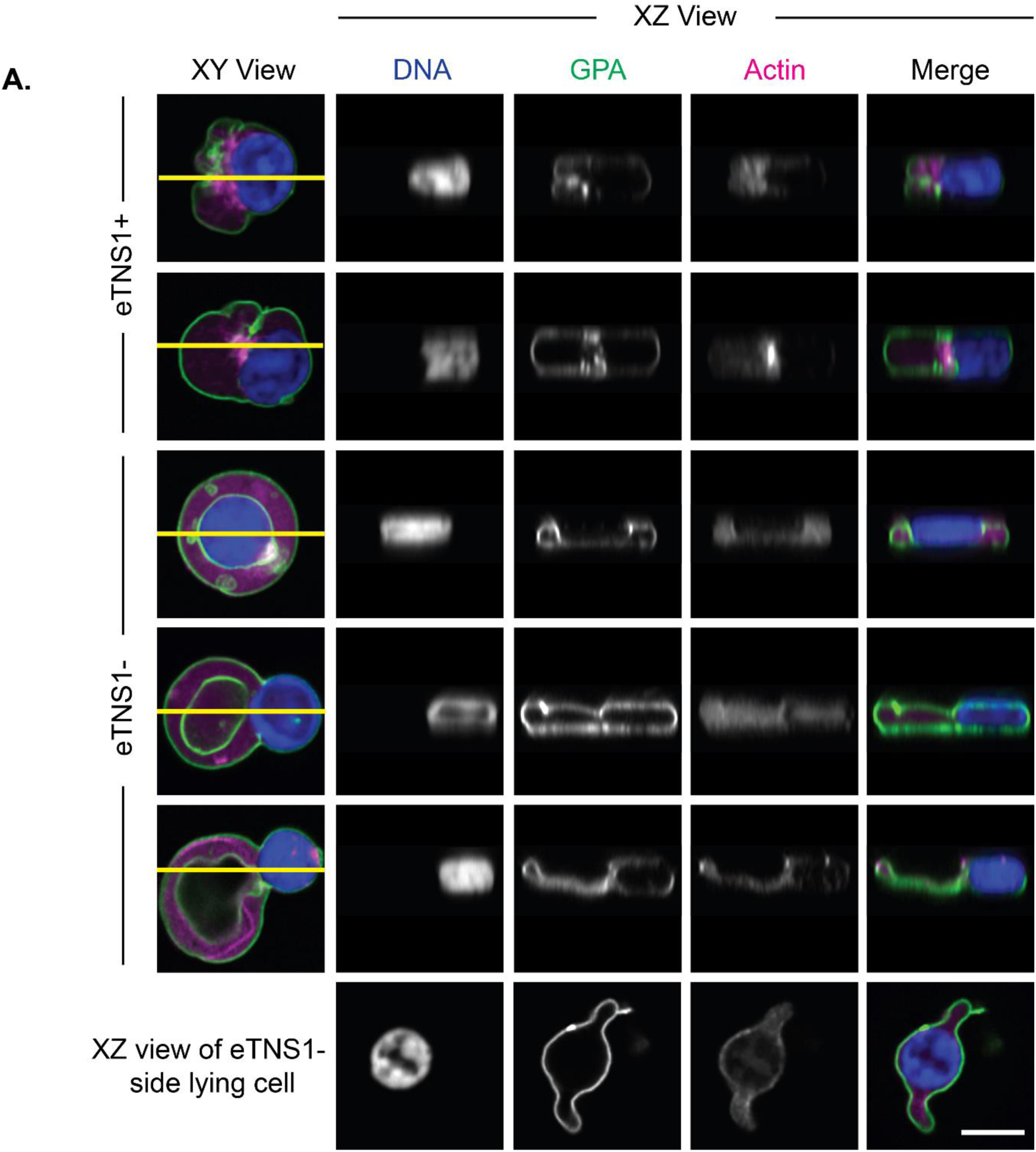
CRISPR/Cas9 knockout of eTNS1 results in human erythroblasts stuck in process of nuclear expulsion, with altered morphology resembling a biconcave cell. (A) Single optical sections from Airyscan Z-stacks of polarized and enucleating eTNS1+ and eTNS1- erythroblasts. eTNS1- cell shapes appeared to resemble biconcave discs (GPA in white), despite the retention of the nucleus in the cell. Z-sections of cells were used to construct an XY side view of each cell at the location of the yellow line. Scale bar, 5 µm.

**Supplemental Table 1: List of known ABPs and NFs used in data mining strategy to identify ABPs/NFs increased during terminal erythroid differentiation**. (A) 135 known ABPs and NFs were identified from multiple studies and compiled into groups based on gene description and actin-associated function. Associated Genbank IDs are provided. Database mining was conducted by comparing this curated list of 135 ABPs/NFs to RNA-seq and proteomics (LC-MS) datasets. (B) For reference controls, 12 known ABPs associated with the red cell membrane skeleton and 25 red cell transmembrane and membrane-associated proteins were identified and were also catalogued for analysis. Data mining of our curated list was benchmarked via comparing the expression profiles of these known red cell membrane ABPs and other red cell membrane proteins.

**Supplemental Table 2: RNA-seq and LC-M/S profiles of compiled ABPs and NFs.** Database mining was conducted by comparing a curated ABP/NF gene list to RNA-seq and LC- M/S hits from Yan et al, 2018 and Gautier et al, 2016 respectively. (A) RNA-seq hits were converted to linear scale for 97 ABP/NFs. The BasoE and OrthoE expression levels were normalized to ProE as shown. (B) RNA-seq hits were also analyzed for expression at the CB, BFU-E and CFU-E stage. The 97 RNA-seq hits were additionally clustered based on expression (low to high, Cluster 1-4) levels based on the CFU-E stage. TNS1 was found in cluster 1. (C) 49 LC-M/S hits were compiled from data mining and certain hits with missing values (BRK1, WIPF1, FMNL1, SPTBN2) were not considered for profiling in Figure 1D.

